# Socialization of *Providencia stuartii* enables resistance to environmental insults

**DOI:** 10.1101/2021.12.20.472897

**Authors:** Julie Lopes, Guillaume Tetreau, Kevin Pounot, Mariam El Khatib, Jacques-Philippe Colletier

## Abstract

*Providencia stuartii* is a highly-social pathogen responsible for nosocomial chronic urinary tract infections. The bacterium indeed forms floating communities of cells (FCC) besides and prior-to canonical surface-attached biofilms (SAB). Within *P. stuartii* FCC, cells are riveted one to another owing to by self-interactions between its porins, viz. Omp-Pst1 and Omp-Pst2. In pathophysiological conditions, *P. stuartii* is principally exposed to high concentrations of urea, ammonia, bicarbonate, creatinine and to large variations of pH, questioning how these environmental cues affect socialization, and whether formation of SAB and FCC protects cells against those. Results from our investigations indicate that FCC and SAB can both form in the urinary tract, endowing cells with increased resistance and fitness. They additionally show that while Omp-Pst1 is the main gateway allowing penetration of urea, bicarbonate and ammonia into the periplasm, expression of Omp-Pst2 enables resistance to them.

## Introduction

Bacterial biofilms are multicellular communities embedded into a self-produced extracellular matrix (ECM) that allows their attachment onto a variety of surfaces^1,2^. The formation of surface-attached biofilms (SAB) enables bacteria to endure environmental insults and to survive in ecological niches otherwise hostile for their planktonic counterparts^1,3–5^. This explains that SAB are found in numerous contrasting environments, ranging from glacial surfaces to ships and cooling systems to the surface of all earth oceans^6,7^. Most worryingly, SAB can niche into human tissues or on prosthetic implants, thereby posing a direct threat to human health. Indeed, SAB-engulfed bacteria exhibit an increased tolerance to antimicrobials agents and to the immune system, making them difficult to eradicate once settled^8^. Accordingly, SAB – and possibly other bacterial socialization modes – are involved in most chronic bacterial infections^1,9,10^.

*Providencia stuartii* is a Gram-negative biofilm-forming opportunistic-pathogen from the *Enterobacteriacae* family, and it is responsible for ~10% of hospital-acquired urinary tract infections. The bacterium is not epidemic, and generally infects immunocompromised individuals such as intensive care unit (ICU) residents and patients under long-term catheterization^11–15^. It is yet highly endemic due to its ability to form biofilms and to its strong intrinsic multidrug resistance (MDR) phenotype. The latter is mostly due to the presence of an inducible chromosomally-encoded AmpC β-lactamase, specifically targeting most penicillins and several cephalosporin antibiotics^16^, but can be further aggravated by horizontal transfer of plasmid-encoded extended-spectrum β-lactamases (ESBLs) and metallo-β-lactamases yielding strains capable of resisting to most β-lactam and carbapenem antibiotics, and therefore leaving only few alternatives for antibiotic treatment^17–20^.

SAB formation has been extensively studied in biofilm-forming human pathogens such as *Bacillus subtilis, Pseudomonas aeruginosa* and *Staphylococcus aureus*^21^, allowing to derive a four-step model. Briefly, planktonic [isolated] bacteria first adhere to a surface, forming a monolayer of cells that further develops into a multi-layer colony upon bacterial division and migration. The multilayered colony then maturates by synthesizing its ECM. Upon exhaustion of nutritional resources, the biofilm releases planktonic cells enabling colonization of other niches^21^. Recently, we showed that *P. stuartii* exploits an additional means of socialization before adhesion of cells onto surfaces, whereby self-matching interactions between the extracellular loops of outer membrane (OM) embedded general-diffusion porins enables formation of floating communities of tightly-packed cells (FCC). Formation of FCC precedes that of SAB, suggesting that the latter form from the sedimentation of the former^22,23^.

Porins are water-filled channels present in the OM of Gram-negative bacteria, whose main function is to ensure influx of hydrophilic solutes and ions into the periplasm. General-diffusion porins (hereafter referred to as porins), which only discriminate solutes on the basis of size and charge, are the most abundantly expressed proteins in the OM, with up to 100,000 copies per cell^24–26^. Down-regulation of porins, effectively reducing the uptake of hydrophilic antibiotics such as β-lactams, was observed to correlate with elicitation of resistance in clinical isolates of *Klebsiella pneunomoniae* and *P. aeruginosa*^25,27–29^. In *P. stuartii* ATCC 29914 (thereafter referred to as *P. stuartii),* two porins are expressed, Omp-Pst1 and Omp-Pst2. Both play a major structural role in *P. stuartii* socialization, enabling the scaffolding of FCC through formation of intercellular dimers of OM-embedded trimers (DOTs) acting as rivets between adjacent cells^30,31^. Omp-Pst1 is ~10 times more expressed than Omp-Pst2 under normal growth conditions, constituting the principal entry route for solutes into the periplasm^23,32^ but also the principal DOT provider. Present at ~10-fold lower concentration, Omp-Pst2 presents a slightly higher propensity to form DOTs (dissociation constant of 0.4 μM compared to 0.6 μM for Omp-Pst1^23^). It was shown that Omp-Pst2 plays a role in the early stages of *P. stuartii* growth, as well as in the adaptation of the bacterium to high concentrations of urea and varying pH^22^. A *P. stuartii* strain knocked-out for Omp-Pst2 could be obtained, but not one for Omp-Pst1, suggesting that only Omp-Pst1 is essential.

The principal pathophysiological habitat of *P. stuartii* in humans is the urinary tract and, accordingly, the bacterium can survive in urine. Urea is the main catabolite present in urine (170 mM), followed by ammonia (25 mM), bicarbonate (25 mM), creatinine (7 mM), magnesium (2.5 mM) and calcium (2 mM)^33,34^. The pH of urine is generally slightly acidic, but it can vary from 6 to 8 depending on the dietary habits of patients and/or their genetic background^34,35^. It was shown that the presence of urea and ions (Ca^2+^ and Mg^2+^) at concentrations similar to those found in the urinary tract does not impact the survival of *P. stuartii*^22^, nor does variation of pH across the pH 6 to pH 9 range. It remains unclear, however, how these environmental cues affect the formation of FCC and SAB, and whether or not socialization into such communities endows *P. stuartii* cells with higher resistance to the cues. These are the questions we sought to provide tentative answers to, in the present article.

To this end, we used a combination of epifluorescence microscopy, dynamic light scattering (DLS), and transcriptomic and proteomic approaches, allowing to investigate the impact of different concentrations of urea, ammonia, bicarbonate, and creatinine on the structuration, fitness and porin content of *P. stuartii* FCC and SAB. We also investigated the impact of sudden variations of pH, such as those that occur along the day of even healthy individuals. Aware of the role played by *P. stuartii* porins in the scaffolding and genesis of FCC and SAB, we paralleled phenotypic investigations with both *in vivo* monitoring of Omp-Pst1 and Omp-Pst2 expression and *in vitro* measurements of their ability to self-associate into DOTs. Our results show the respective implication of Omp-Pst1 and Omp-Pst2 in the uptake-of and resistance-to harmful solutes, and establish that socialization into FCC increases the tolerance of *P. stuartii* cells to a variety of chemical aggressions. Thus, atop providing new insights into the formation and resistance of *P. stuartii* communities in their pathophysiological environment, our results suggest that future treatments against the bacterium should be tested on FCC and SAB, and in conditions mimicking the urinary tract; both supracellular structures are indeed able to form and survive in this habitat.

## Results

We used reverse transcription quantitative PCR (RT-qPCR) to monitor the expression of the genes encoding the two porins of *P. stuartii* ATCC29914, *omp-pst1* and *omp-pst2,* at different stages of growth and in both FCC (Figure 1.a) and SAB (Figure 1.b). FCC develop after 4 hours of growth and sediment ~3 hours later (7h) to form SAB. Regardless of the stage of growth or of the socialization mode, we found that Omp-Pst1 is 10-15 fold more expressed than Omp-Pst2, in line with previous results^23^ and with the proposition that Omp-Pst1 is the major porin in *P. stuartii*^22,32^. In both phenotypes, a comparable level of expression was observed at all stages, i.e. from 4 to 15 h post-inoculation for FCC, and 7 to 15 h post-inoculation in SAB, respectively.

**Figure 1.**
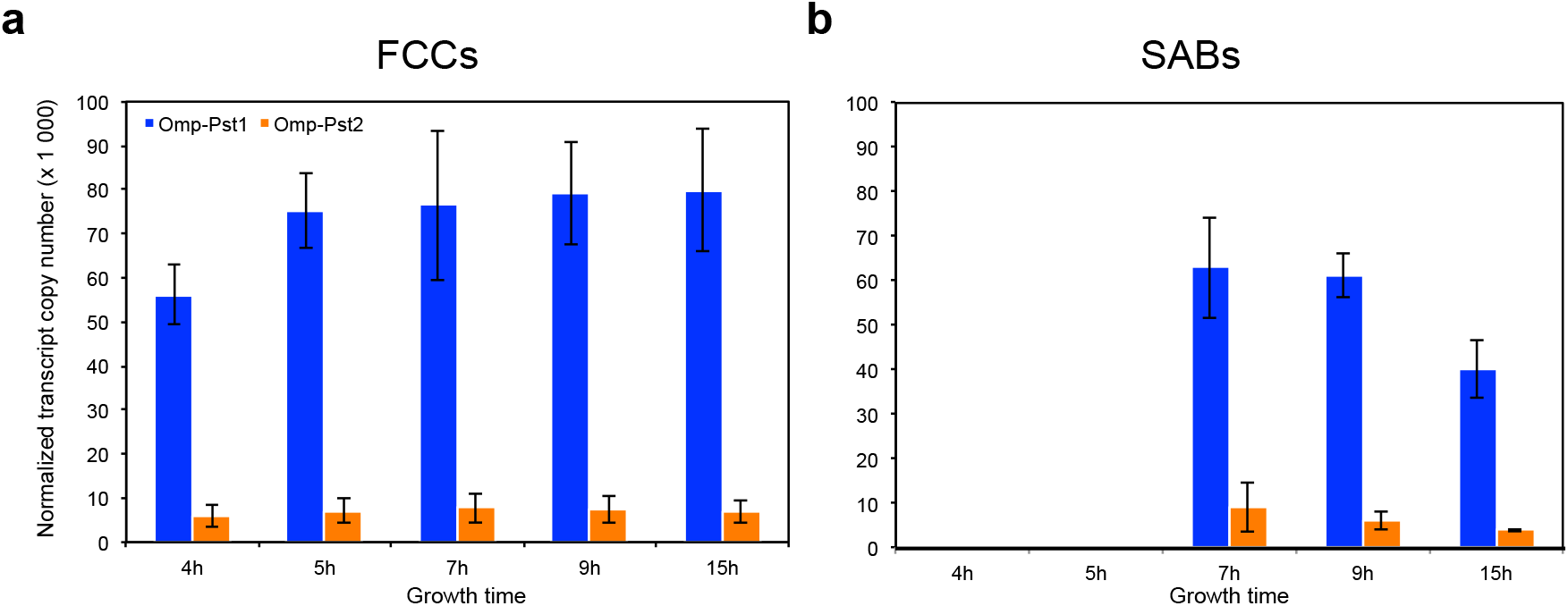
Omp-Pst1 is the most abundantly expressed porin in P. stuartii, regardless of the socialization mode. Expression of *omp-pst1* (blue) and *omp-pst2* (orange) porins was monitored by RT-qPCR, at different stages of growth, and both in FCC (panel a) and SAB (b). Reported numbers of normalized transcript copy were each averaged over three biological replicates; errors bars outline the standard deviation of measurements.

### P. stuartii cells are highly resistant to urea

The fitness and survival of *P. stuartii* cells in the presence of increasing concentrations of urea were assessed by optical density (OD) measurements and epifluorescence microscopy (Figure 2 and Supplementary Figure S1). No effect was observed on bacterial growth and socialization at concentrations up to 150 mM urea. At 500 mM urea, the reach of the exponential phase was delayed by 5 hours, yet neither the formation of FCC nor SAB was affected. At 1,000 mM urea, no bacterial growth was observed, indicating that the mechanism enabling adaptation to high urea concentration saturates above 500 mM (Figure 2).

**Figure 2.**
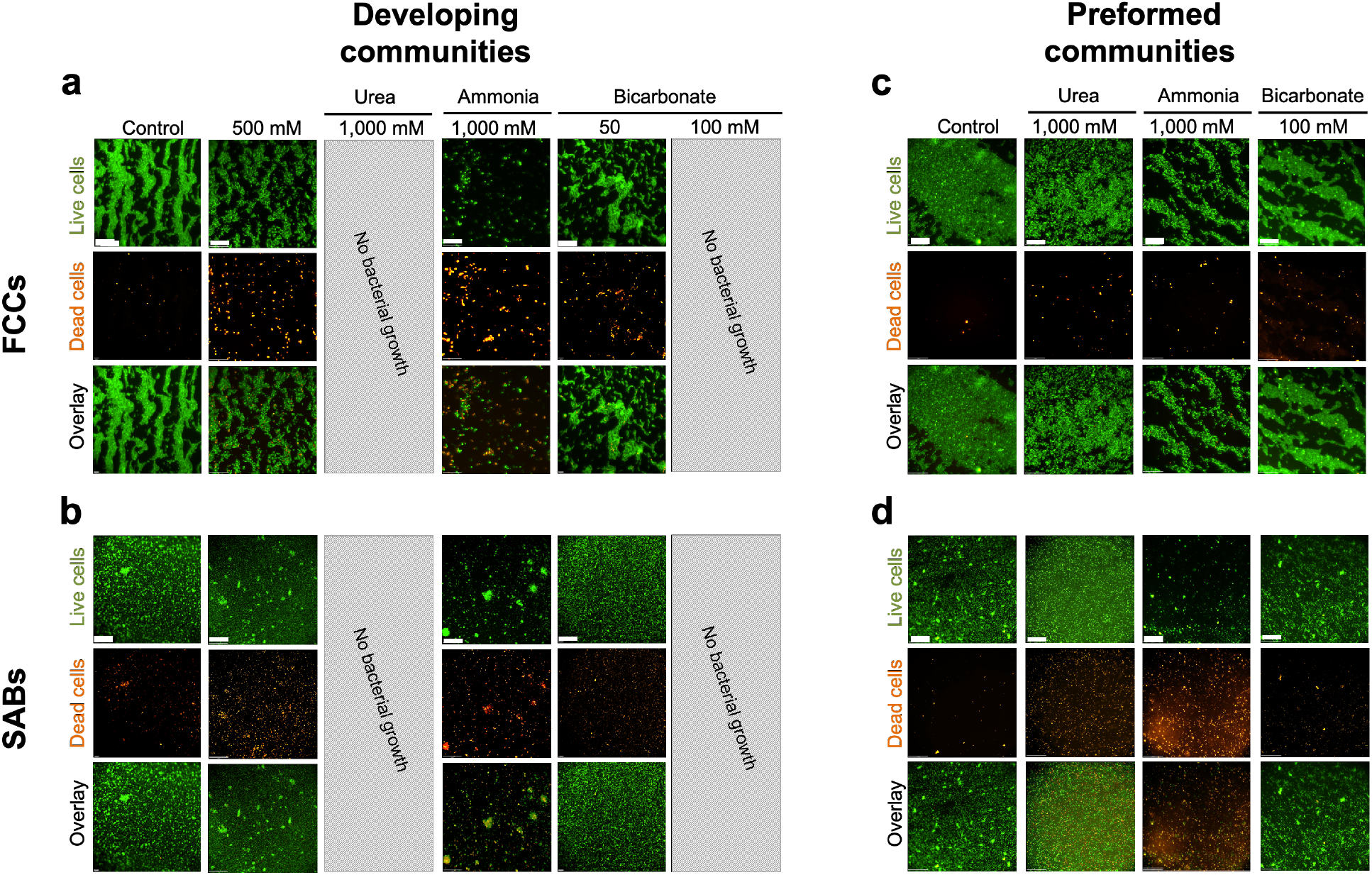
Effect of urea, ammonia and bicarbonate on developing and preformed FCC and SAB. Epifluorescence microscopy was used to monitor FCC and SAB formation (‘developing communities’) and survival (‘preformed communities’) in presence of various cues. All bacteria are labelled by the permeant DNA stain Syto9 (green channel), but only dead cells are labelled by propidium iodide (red channel). For each condition, an overlay of the two channels is shown. Grey-hatched rectangles indicate concentrations at which bacteria were unable to grow.

Porins being the main gateway for hydrophilic solutes into Gram-negative cells, we examined whether exposure to increasing urea concentrations affects their expression. At the proteomic level, *i.e.* as assessed from the intensity of bands on SDS-PAGE gels of OM extracts (see Methods), we found that urea concentrations up to 150 mM have no effect on porin abundance, while a 500 mM concentration results in a 33% decrease in the overall abundance of porins in the OM (Supplementary Figure S2 and Table S1). This concentration was therefore chosen to further investigate the effect of urea on porin genes transcription in growing FCC and SAB (Supplementary Figure S2). Using RT-qPCR, we found that in the early stages of growth (~4 h post-inoculation), i.e. when only FCC are present, the presence of 500 mM urea results in a 4-fold over-expression of Omp-Pst2. In contrast, the expression of Omp-Pst1 remains unchanged (Figure 3.a and Supplementary Table S2). In SAB, which form from the sedimentation of FCC after ~7 hours of growth, neither the expression of Omp-Pst1 nor of Omp-Pst2 is significantly affected by the presence of urea (Figure 3.b and Supplementary Table S2).

**Figure 3.**
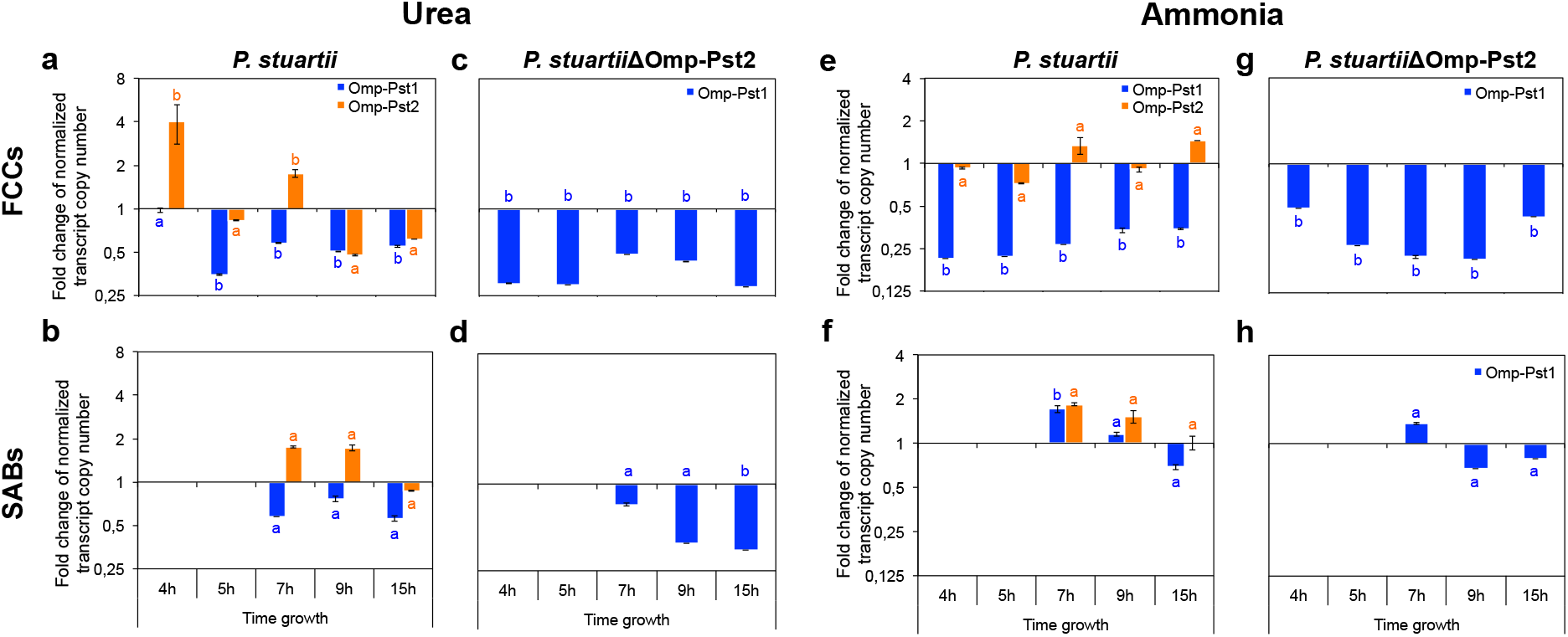
Presence of urea or ammonia induces a down-regulation of porin expression in P. stuartii. Expression of *omp-pst1* (blue) and *omp-pst2* (orange) porins in presence of 500 mM urea (a,b,c,d) or 500 mM ammonia (e,f,g,h) was monitored by RT-qPCR in both *P. stuartii* (a,b,e,f) or *P. stuartii*ΔOmp-Pst2 (c,d,g,h), at different stages of growth, and both in FCC (panel a,c,e,g) and SAB (b,d,f,h). For each growth time point, we report the ratio of normalized transcript copy numbers between cells exposed and not exposed (control) to the cues. All measurements were performed in triplicates; errors bars outline standard deviations from the average. Different letters above the bars indicate significant differences (p < 0.05; ANOVA followed by post-hoc Tukey HSD test; see statistical indicators in Supplementary Table S2).

To characterize the effect of urea on already socialized *P. stuartii* cells, FCC and SAB were grown in the absence of urea, and then exposed to increasing concentrations of the catabolite. FCC and SAB formation were monitored by epifluorescence microscopy (Figure 2 and Supplementary Figure S3), and the ability of socialized bacteria to resume growth by OD measurements. Cells were not affected by exposure to 150 mM urea but lag times of 2 and 2.5 hours were observed in the presence of 500 and 1,000 mM of urea, respectively. Regardless, socialization was not affected by urea, with FCC and SAB maintaining their organization and morphology even at urea concentrations as high as 1,000 mM. This observation suggests that once formed, FCC and SAB can endure urea concentrations higher than they would have in their early stage of growth.

To shed further light on the possible role of Omp-Pst2 in the adaptation to high urea concentration, we repeated our experiments on a strain of *P. stuartii* knocked-out for this porin (called hereafter *P. stuartii*ΔOmp-Pst2), i.e. expressing only Omp-Pst1 as a general diffusion porin. This strain retains the ability of the parental wild-type (WT) strain to socialize into FCC and SAB but is characterized by a 6-hours lag-time in normal (laboratory) growth conditions^22^. Absence of Omp-Pst2 resulted in a lag time of ~10 hours in presence of 150 mM urea, again indicating an important role for this porin in the early stages of growth (Supplementary Figure S4). The capacity of cells to form FCC and SAB was however not impacted, nor was their capacity to endure the presence of urea at up to 500 mM concentration (Supplementary Figure S4). Note that the expression of Omp-Pst1 is 3-fold down-regulated within developing *P. stuartii*ΔOmp-Pst2 FCC and SAB at this concentration (Figure 3.c and d, respectively, and Supplementary Table S2). Thus, Omp-Pst2 could play a role in the resistance of *P. stuartii* communities to high urea concentrations – a role that could be diffusive (efflux or reduced influx of urea) or structural (promotion of DOT formation). To determine whether urea influences the formation of DOT by Omp-Pst1 and Omp-Pst2, we reconstituted these in liposomes and assayed by use of dynamic light scattering (DLS), the porin-induced proteoliposome aggregation, characteristic of DOT formation^23^, at increasing concentrations of urea (Supplementary Figure S5 and Table S3). We found that urea does not affect the self-association into DOTs of *P. stuartii* porins, consistent with the observation that urea does not impact FCC formation.

### Resistance of P. stuartii to high concentration of ammonia involves Omp-Pst2

We investigated the effect of high concentrations of ammonia on *P. stuartii* fitness, survival, socialization, and on porin expression and self-association using the same approaches described above for urea. At the growth level, the presence of ammonia at 500 and 1,000 mM concentrations resulted in lag times of 45 min and 6.4 hours, respectively, with no effect seen at 150 mM (Figure 2 and Supplementary Figure S1). The increased lag times were paralleled by significant decreases in the abundance of porins in the OM, viz. - 20 and - 50% at 500 and 1,000 mM ammonia, respectively (Supplementary Figure S2 and Table S1). Transcriptomic studies revealed that only expression of Omp-Pst1 is affected, with a significant 4-fold reduction in expression observed at 500 mM ammonia in developing FCC (Figure 3.e and Supplementary Table S2). Contrastingly, Omp-Pst1 expression is 2-fold increased in developing SAB exposed to the same concentration of ammonia (Figure 3.f and Supplementary Table S2). Within both type of communities, the expression of Omp-Pst2 remains steady (Figure 3.e and f). Thus, the regulation of Omp-Pst1 expression in the presence of ammonia is different in SAB and FCC, possibly underlying differences in the access of this solute to the periplasm in the two types of communities. Indeed, an extracellular matrix (ECM) is present around SAB cells which could protect them against the adverse effects of ammonia. As this matrix is not visible and therefore presumably absent in FCC, reduction of Omp-Pst1 expression could in these serve the purpose of reducing penetration of the ammonia into the periplasm. Conversely, the observation that the expression of Omp-Pst2 is not down-regulated suggests that it does not partake in ammonia influx. This hypothesis is supported by experiments performed on the *P. stuartii*ΔOmp-Pst2 strain which confirm that a significant down-regulation of Omp-Pst1 expression (~ 2-4 fold) is required for FCC to survive at high ammonia concentration (Figure 3.g and Supplementary Table S2). The observation that deletion of Omp-Pst2 results in an inability to grow at 1,000 mM ammonia furthermore suggests that expression of Omp-Pst2 is beneficial to *P. stuartii* survival and fitness when this solute is present (Supplementary Figure S4).

Socialization was challenged by high concentrations of ammonia, with a complete inability to form SAB at 1,000 mM ammonia. Exposure of preformed FCC to high concentrations of ammonia (≥ 500 mM) results in their breaking into smaller communities of living cells, whereas preformed SAB are disrupted at 1,000 mM ammonia with only dead cells remaining attached onto the surface (Figure 2.a and b). This result evidences that for an ECM to efficiently protect cells against ammonia, it needs to be produced in the presence of ammonia. It also demonstrates that formation of FCC can in some casesbe a more fit socialization mechanism for *P. stuartii* than SAB, e.g. in the presence of ammonia. The observation that socialization is not affected by the deletion of Omp-Pst2 supports the hypothesis that Omp-Pst2 does not partake in the influx of ammonia into the periplasm (Supplementary Figure S4). It also exemplifies that in the presence of ammonia, Omp-Pst1 is the main porin supporting cell-to-cell contact by DOT formation in the FCC.

We asked whether the formation of smaller FCC in the presence of high ammonia concentrations only parallels the observed reduction in porin expression or is associated with a reduced propensity of porins to self-associate. We found that ammonia significantly inhibits DOT formation by Omp-Pst1, but not that by Omp-Pst2 (Figure 4.a and Supplementary Table S3). Indeed, the proteoliposome aggregation driven by 1 μM Omp-Pst1 can be fully reversed by concentration as low as 150 mM ammonia, whereas that of Omp-Pst2 is not affected even at concentration as high as 1,000 mM. Thus, the formation of smaller FCC stems both from the reduced number of Omp-Pst1 present in the OM and from the inhibition of Omp-Pst1 DOT at high ammonia concentration.

**Figure 4.**
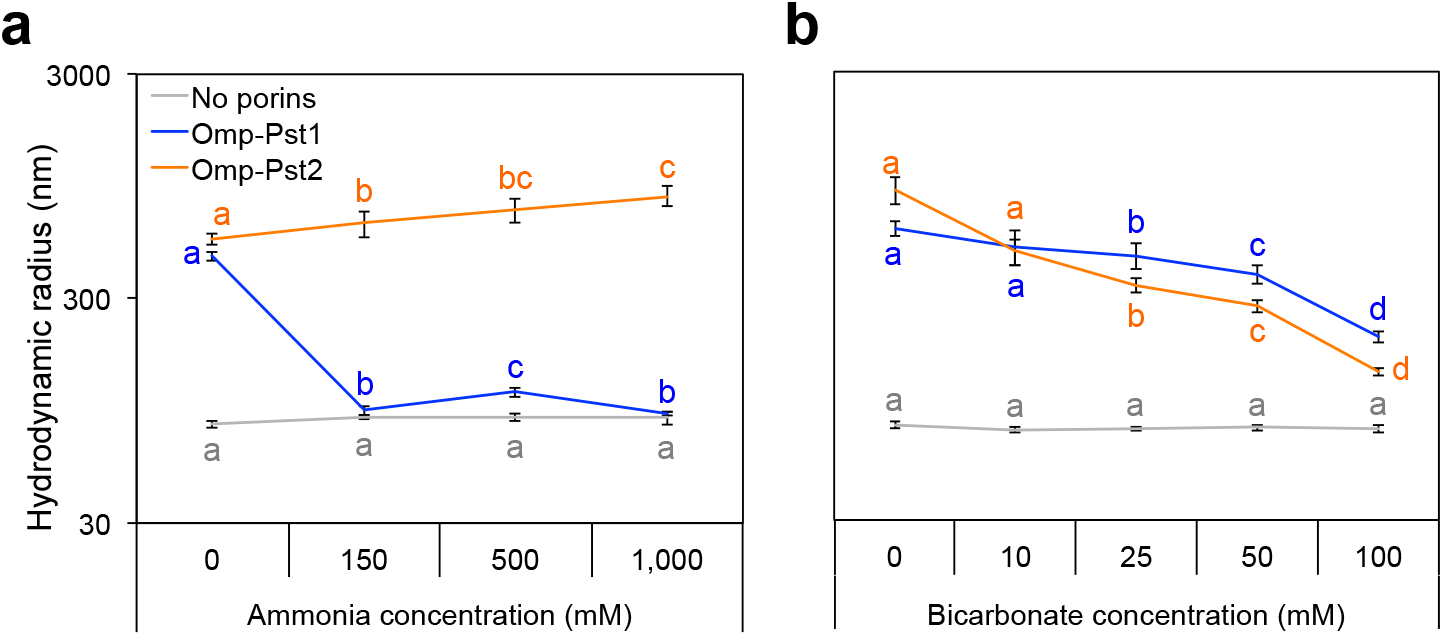
Ammonium and bicarbonate both inhibit self-association of Omp-Pst1, but that of Omp-Pst2 is only challenged by bicarbonate. LDAO-solubilized Omp-Pst1 (blue) and Omp-Pst2 (orange) were reconstituted in ~50-nm radius LUV, in presence of increasing concentrations of ammonia (a) or bicarbonate (b), and after 24 h incubation with biobeads, the hydrodynamic radius of proteoliposomes was measured by DLS. The hydrodynamic radii of LUVs incubated at with the cues but without porin were also measured (grey). Different letters above the bars indicate significant differences (p < 0.05; ANOVA followed by post-hoc Tukey HSD test; see statistical indicators in Supplementary Table S3).

### P. stuartii is highly sensitive to bicarbonate but unaffected by creatinine

Bicarbonate and creatinine are the two most abundant metabolites present in the urine, after urea and ammonia. Performing the same investigations as described above for urea and ammonia, we found that up to 100 mM, i.e. a concentration 14-time higher than encountered in the urinary tract (7 mM), creatinine has no effect on *P. stuartii*fitness, survival, socialization (Supplementary Figure S1 and S4) and porin self-association (Supplementary Figure S5).

*P. stuartii* cells are, however, very sensitive to changes in bicarbonate concentration, displaying all signs of normal growth at 50 mM (Figure 2 and Supplementary Figure S1), but a complete inability to develop FCC and SAB at 100 mM (Figure 2.c and d, respectively). Moreover, a decrease in size of the FCC is observed when the bicarbonate concentration is increased (up to 50 mM), coherent with the inhibition of Omp-Pst1 and Omp-Pst2 self-association into DOTs (Figure 4.b and Supplementary Table S3). However, full inhibition of DOT formation is not attained at 100 mM bicarbonate, supporting that the observed inhibition of *P. stuartii* socialization is not only due to inhibition of porin self-association.

Accordingly, RT-qPCR indicates that Omp-Pst1 undergoes a 2-fold down-regulation in expression just post-exposure to bicarbonate at 50 mM concentration, although we note that expression of Omp-Pst1 is restored in FCC along time, and even augmented in SAB. Contrastingly, the expression of Omp-Pst2 is slightly augmented in WT FCC (Supplementary Figure S2 and Supplementary Table S3) while deletion of this porin in the *P. stuartii* ΔOmp-Pst2 strain renders cells more sensitive to bicarbonate, with growth being observed up to 25 mM bicarbonate only (Supplementary Figure S4). Thus, Omp-Pst2 is necessary for *P. stuartii* cells to survive in the presence of bicarbonate. The expression of neither porin varies significantly in SAB, again supporting that the presence of an ECM around SAB cells enables resistance to bicarbonate without necessity of porin regulation.

We then assayed the effect of bicarbonate on already established communities of *P. stuartii* cells and found that both preformed FCC and SAB are able to resume growth without delay at up to 100 mM bicarbonate (Supplementary Figure S3). These results again illustrate that socialization profits *P. stuartii* survival. They also raise questions as to which mechanism(s) Omp-Pst2 may exploit to protect cells against the adverse effects of bicarbonate.

### Variation of pH hardly affects P. stuartii socialization and fitness

We last investigated the effect of pH variation. As reported earlier, *P. stuartii* can grow and socialize into FCC and SAB within the pH 6 to pH 9 range^22^. Within FCC, however, growth at extreme pHs is poised with increased lag times (5, 1 and 2 hours lag time at pH 5, 8 and pH 9, respectively) (Supplementary Figure S1) and characterized by a 20% reduction in the abundance of porins in the OM at pH 9 (Supplementary Figure S2 and Table S1). RT-qPCR experiments indicate that this decrease can be fully accounted-for by Omp-Pst1, whose expression is down-regulated by 2.6-fold in FCC at pH 9, while that of Omp-Pst2 remains steady. Within SAB, the situation is different and dependent on pH, with a 2.1-fold overexpression at pH 5 but at 1.6-fold reduced expression at pH 9 (Supplementary Figure S2 and Table S2). Note that no change in porin expression is observed along time, regardless of the socialization mechanism (Supplementary Table S2).

Experiments performed on the *P. stuartii*ΔOmp-Pst2 strain suggest that Omp-Pst2 plays a role in the adaptation of cells to variations of pH. Firstly, lag times before growth are increased at pH 5 and pH 9 (6 and 7 hours lag time, respectively) and, to a lesser extent, at pH 8 (~3 hours lag time), in the absence of Omp-Pst2 (Supplementary Figure S4). Secondly, a significant increase is observed in the amount of Omp-Pst1 present within OM at pH 8 and 9 (Supplementary Figure S2 and Table S1), suggesting that *P. stuartii* compensates the absence of Omp-Pst2 by increasing the amount of Omp-Pst1. Unfortunately, we cannot draw a parallel with transcriptomic results due to high variations in Omp-Pst1 expression within FCC and SAB, over the time. We note that while variations of pH do not affect the propensity of Omp-Pst2 to self-associate into DOTs, that of Omp-Pst1 decreases by 2.1 and 2.3 times at pH 8 and pH 9, respectively (Supplementary Figure S5 and Table S3).

## Discussion

*P. stuarti* is, of all strains in the *Enterobacteriaceae family,* that displaying the most stringent MDR phenotype. It is also amidst the most social bacterial strains, displaying the ability to socialize both into floating communities of cells (FCC) and surface-attached biofilms (SAB)^23^. These facts together explain that urinary tract infections by *P. stuartii* are generally chronic and can result in the death of victims^11,15^.

FCC are supported by self-matching interaction between residues in the extracellular loops of porins, yielding DOTs able to rivet cells across. FCC are at the origin of SAB, which they form by sedimentation^23^ and subsequent secretion of an extracellular matrix (ECM). We examined the impact of a variety of environmental cues on the fitness, survival and socialization of *P. stuartii,* as well as on the expression of its porins and their propensity to self-associate into DOTs. Experiments were hence performed on both types of multicellular communities (FCC and SAB), and on both developing and established communities. The effect of four metabolites present at high concentration in the urinary tract was investigated, viz. urea (up to 170 mM), ammonia (25 mM), bicarbonate (25 mM) and creatinine (7 mM); we also verified the effect of sudden pH variations. Overall, we found that *P. stuartii* is highly resistant to these cues, requiring 2-40 times higher concentrations than those encountered in the urinary tract to display a phenotypic change. Socialization into FCC and SAB was observed in presence of all tested environmental cues and at all concentrations tested, with the exception of bicarbonate at 100 mM. Given that this concentration is 4-times that observed for bicarbonate in the urinary tract, our results are indicative of the bacterium likely being present in a socialized state (FCC and SAB) in the urinary tract, during infections. Comparison of results obtained on developing and preformed FCC and SAB suggests that socialization increases the fitness of *P. stuartii,* raising tolerated concentrations by at least 2-fold irrespective of the tested cue. For example, bicarbonate is the most harmful to *P. stuartii* cells with no growth observed at a concentration as low as 50 mM, but preformed FCC and SAB can survive up to 100 mM concentration. Likewise, urea is the second most toxic metabolite we tested, with no growth observed at 1,000 mM concentration, yet preformed FCC and SAB can sustain this concentration. In the case of ammonia, which is less toxic to *P. stuartii* cells with growth being observed up to 1,000 mM concentration, the increased fitness of socialized cells was visible in the lag times before growth, those being 1.5 and 6.4 h in preformed and developing FCC, respectively. Thus FCC represent a socialization mode which, alike SAB, benefits cells by allowing them to endure higher concentrations of harmful solutes.

Experiments performed on the *P. stuartii*ΔOmp-Pst2 strain show that DOTs of Omp-Pst1 are sufficient to scaffold FCC, as one would expect given the relatively high abundance of Omp-Pst1 in *P. stuartii* OM, as compared to Omp-Pst2. They also demonstrate that the presence of Omp-Pst2 is beneficial to *P. stuartii* fitness, enabling survival at 2-times higher ammonia and bicarbonate concentrations (1,000 & 500 mM ammonia and 50 & 25 mM bicarbonate for *P. stuartii* WT & *P. stuartiiΔOmp-Pst2,* respectively), and reducing lag-time before growth at high concentrations of urea and ammonia (5 & 9 hours at 500 mM urea and 6.5 hours & no growth at 1,000 mM ammonia for WT & *P. stuartiiΔOmp-Pst2,* respectively). At the present time, we can only speculate as to how Omp-Pst2 promotes resistance against high urea, ammonia and bicarbonate concentrations. Alike its role in the early stage of growth, it could either be diffusive, i.e. a result of the peculiar positively-charged amino-acid distribution along the channel of Omp-Pst2^32,36,37^, or structural, i.e. stemming from the ability of this porin to form intercellular dimers of trimers (DOTs), riveting adjacent cells one onto another across FCC^23^. We conjecture that it is unlikely that influx of urea, ammonia or bicarbonate occurs across Omp-Pst2, whose electrostatic potential favors the efflux of cations and gates the channel in case of a massive cation influx^36^. Rather, our data suggest that, regardless of whether Omp-Pst2 is expressed (*P. stuartii* WT) or not (*P. stuartii*ΔOmp-Pst2), Omp-Pst1 serves as the entry route for these solutes into cells, with a ~2-4 fold reduced expression in FCC and SAB formed in presence of 500 mM urea, 500 mM ammonia or 50 mM bicarbonate. Hence, if any, the diffusive role of Omp-Pst2 in resistance to high ammonia concentrations would be to facilitate the efflux of positively charged ammonia from the periplasm^36^, following urea catalysis by the cytoplasmic urease of *P. stuartii*or degradation by selfhydrolysis. The role of Omp-Pst2 could alternatively be to provide a substitute scaffold to promote FCC formation when expression of Omp-Pst1 needs to be repressed. In line with this hypothesis, the propensity of Omp-Pst2 to self-associate into DOTs is slightly higher than that of Omp-Pst1^23^ and is promoted by increasing ammonia concentrations. Together with the fact that DOT formation by Omp-Pst1 and Omp-Pst2 are unaffected by urea, this compensation mechanism could explain why cells exposed to 500 mM urea are still able to form FCC despite a 33% reduced abundance of porin in their OM. Close-up views of these FCC indeed show the characteristic tight packing and “standing” orientation of cells – almost orthogonal to that observed in micro-colonies (Supplementary Figure S6) – confirming that even at high urea concentration (and therefore low cell density), the structural integrity of FCC is preserved. To the contrary, FCC formed at 500 mM ammonia show defects with cells losing their “standing” orientation (Supplementary Figure S6), possibly as a combined consequence of the 50% reduction in porin abundance in the OM, accounted entirely by Omp-Pst1 (no change in Omp-Pst2 expression in the *P. stuartii* WT strain and same expression profile for Omp-Pst1 in the *P. stuartii*ΔOmp-Pst2 strain), and the marked inhibition of Omp-Pst1 self-association by ammonia. Again, Omp-Pst2 could compensate – whose self-association into DOTs is promoted by increasing ammonia concentrations – yet visibly not up to 1,000 mM ammonia, where planktonic cells are observed instead of FCC. Loss of the ability to scaffold FCC and/or to efflux ammonia would then explain why deletion of Omp-Pst2 result in an increased sensitivity to ammonia. By extension, bicarbonate influx likely occurs through Omp-Pst1 – whose channel is mildly anion selective – considering the immediate down-regulation undergone by this porin when the bacterium is put in presence of the solute. The increase in Omp-Pst2 expression was yet observed only after 15 hours, suggesting that it does not come into play to rescue FCC scaffolding by providing an alternative DOT architecture. In line with this premise, bicarbonate displayed an inhibitory effect on DOT formation by both porins, likely explaining the disruption of FCC into planktonic cells.

We explained earlier that FCC are at the origin of SAB, which they form by sedimentation. SAB are characterized by an ECM, but it is unknown whether such a structure exists around FCC, or in other words whether synthesis of the ECM debuts or not at the FCC stage. In the present work, we have tried to specifically label the ECM of *P. stuartii* using three common markers (viz. Wheat Germ Agglutinin (WGA) lectin, Concanavaline A lectin and FilmTracer^TM^ SYPRO^TM^ Ruby Biofilm Matrix Strain (ThermoFischer)), none of which succeeded in revealing the ECM of *P. stuartii* SAB; hence the question as to whether or not FCC also feature an ECM remains open. Data collected in the presence of 500 mM ammonia, however, showed a 3-fold reduction in Omp-Pst1 expression in developing FCC, whereas a 2-fold increase in the expression of the two porins was observed in developing SAB. This difference could underlie the fact that in the latter an ECM is present that protects cells against the adverse effects of ammonia, eliminating the need to down-regulate porin expression. Our results yet evidence that the ECM may only be able to do so in the case it has been synthesized in the presence of ammonia; indeed SAB preformed in normal growth conditions are eradicated by exposure to 1,000 mM ammonia. We note that preformed FCC can sustain this concentration, demonstrating that in the presence of ammonia at least they represent a more fit socialization mechanism than SAB.

In conclusion, our investigations show that FCC and SAB can both form in the presence of urinary tract cues, endowing cells with increased resistance and fitness. They establish a direct link between porins, FCC and SAB, showing that in conditions where porin expression is repressed and their self-association into DOT inhibited, FCC do not form preventing the appearance of SAB. They show that while Omp-Pst1 is the main gateway allowing penetration of urea, bicarbonate and ammonia into the periplasm, expression of Omp-Pst2 enables resistance to them. In the case of ammonia, data show that exposure of cells at the FCC stage is required for them to resist to this cue at the SAB stage, indicating that SAB formed ex-vivo can be eliminated by treatment at high ammonia concentration (1000 mM). They furthermore reveal that bicarbonate can be used as a low-cost cleansing agent capable of eradicating both FCC and SAB at concentration as low as 100 mM. *P. stuartii* is most often isolated from patients undergoing long-term catheterization^11,15^, and the severity of infections results not only from the intrinsic antibiotic resistance of the bacterium, but also from its socialization into FCC and SAB^22^, which display higher resistance to environmental cues (this work), antibiotics and the immune system^38,39^. Our data indicate that future treatments against *P. stuartii* should be evaluated on FCC and SAB, and in conditions mimicking the urinary tract, for them to be efficient and potentially eradicate infections in the clinical context.

**Supplementary Figure S1. *P. stuartii* is highly resistant to catabolites present in the urinary tract. (a-d)** The overall impact of environmental cues on *P. stuartii* growth was monitored by determining, at increasing concentrations of the cues or pH, the lag time before reaching an optical density of 0.2 at 600 nm. Panels a, b, c, and d show results for urea, ammonia, bicarbonate and for different pH, respectively. All measurements were performed in triplicates; errors bars outline standard deviations from the average. Grey-hatched rectangles indicate concentrations at which bacteria were unable to grow. (**e-n**) Epifluorescence microscopy was used to monitor FCC (e,g,I,k,m) and SAB formation (f,h,j,l,n) in the presence of urea (e,f), ammonium (g,h), bicarbonate (i,j), creatinine (k,l) and at various pH (m,n). All bacteria are labelled by the permeant DNA stain Syto9 (green channel), but only dead cells are labelled by propidium iodide (red channel). For each condition, an overlay of the two channels is shown. Grey-hatched rectangles indicate concentrations at which bacteria were unable to grow. Scale bars correspond to 50 and 200 μm in FCC and SAB micrographs, respectively.

**Supplementary Figure S2. Regulation of porin expression in the presence of environmental cues is more pronounced in FCC than SAB, and sometimes opposed. (a-d)** Porin abundance in the OM of *P. stuartii* FCC and SAB cells grown in presence of increasing concentrations of urea (panel a), ammonia (b), bicarbonate (c) and to various pH (d) was evaluated by image processing of digitized SDS-PAGE gels using ImageJ. Plots show percent increase or decrease in porin abundance in the OM, after normalization of intensity counts from porin bands of exposed bacteria by those of unexposed bacteria. All measurements were performed in triplicates; errors bars outline standard deviations from the average. Different letters above the bars indicate significant differences (p < 0.05; ANOVA followed by post-hoc Tukey HSD test; see statistical indicators in Supplementary Table S1). **(e-v)** Immediate changes in the expression of Omp-Pst1 (blue) and Omp-Pst2 (orange) in WT *P. stuartii* or *P. stuartii*ΔOmp-Pst2 FCCs and SABs cells were monitored by RT-qPCR following 30 min incubation with 500 mM of urea (panels e, f and g, h, respectively), 500 mM ammonia (i, j and k, l), to 50 mM bicarbonate (m, n and o, p), or to pH variation (q, r, s, t and u, v). For each growth time point, we report the ratio of normalized transcript copy numbers between cells exposed and not exposed (control) to the cues. All measurements were performed in triplicates; errors bars outline standard deviations from the average. Different letters above the bars indicate significant differences (p < 0.05; ANOVA followed by post-hoc Tukey HSD test; see statistical indicators in Supplementary Table S2).

**Supplementary Figure S3. FCC and SAB cells are generally more resistant than their plantktonic counterparts. (a-b)** Recovery of preformed *P. stuartii* FCC and SAB cells after sudden exposure to urea (a) or ammonia (b) was monitored by determining, at increasing concentrations of these, the lag time before reaching an optical density of 0.2 at 600 nm. All measurements were performed in triplicates; errors bars outline standard deviations from the average. (**e-h**) Epifluorescence microscopy was used to monitor FCC (e,g,I,k,m) and SAB survival and recovery (f,h,j,l,n) in presence of increasing concentrations of urea (panels c and d, respectively), ammonia (e, f), and bicarbonate (g, h). All bacteria are labelled by the permeant DNA stain Syto9 (green channel), but only dead cells are labelled by propidium iodide (red channel). For each condition, an overlay of the two channels is shown. Grey-hatched rectangles indicate concentrations at which bacteria were unable to grow. Scale bars correspond to 50 and 200 μm in FCC and SAB micrographs, respectively. Note that preformed *P. stuartii* FCC and SAB generally resist better to the cues than their developing counterparts, as illustrated by survival of FCC and SAB at 1 M urea and 100 mM bicarbonate, where new cells do no grow (Supplementary Fig. 1). An exception is ammonium, which disrupts SAB formed in its absence although preformed FCC survive and new FCC and SAB can form (Supplementary Fig. 1).

**Supplementary Figure S4. Omp-Pst2 benefits resistance of *P. stuartii* to its pathophysiological environment. (a-d)** The contribution of Omp-Pst2 in adapting to high concentration of urea (a), ammonia (b), bicarbonate (c) or to different pHs (d) was evaluated on the *P. stuartii*ΔOmp-Pst2 strain by determining, at increasing concentrations of the cues or pH, the lag time before reaching an optical density of 0.2 at 600 nm, and comparing results to the wildtype (Supplementary Fig. 1). All measurements were performed in triplicates; errors bars outline standard deviations from the average. Grey-hatched rectangles indicate concentrations at which bacteria were unable to grow. (**e-n**) Epifluorescence microscopy was used to monitor FCC (e,g,I,k,m) and SAB formation (f,h,j,l,n) by *P. stuartii*ΔOmp-Pst2 cells in the presence of urea (e,f), ammonium (g,h), bicarbonate (i,j), creatinine (k,l) and at various pH (m,n). All bacteria are labelled by the permeant DNA stain Syto9 (green channel), but only dead cells are labelled by propidium iodide (red channel). For each condition, an overlay of the two channels is shown. Grey-hatched rectangles indicate concentrations at which bacteria were unable to grow. Scale bars correspond to 50 and 200 μm in FCC and SAB micrographs, respectively.

**Supplementary Figure S5. Omp-Pst1 and Omp-Pst2 self-association into DOT can be challenged by environmental cues.** LDAO-solubilized Omp-Pst1 (blue) and Omp-Pst2 (orange) were reconstituted in ~50-nm radius LUV, in presence of increasing concentrations of urea (a), ammonia (b), bicarbonate (c), creatinine (d) or at various pH (e), and after 24 h incubation with biobeads, the hydrodynamic radius of proteoliposomes was measured by DLS. The hydrodynamic radii of LUVs incubated at with the cues but without porin were also measured (grey). Different letters above the bars indicate significant differences (p < 0.05; ANOVA followed by post-hoc Tukey HSD test; see statistical indicators in Supplementary Table S3).

**Supplementary Figure S6. The orientation of *P. stuartii* cells differs in native FCC and in FCC formed in presence of environmental cues.** Close-up views of FCCs developed in the presence of urea (a) and ammonia (b), showing “standing” and “seated” orientations of cells, respectively. (c) Unexposed FCC feature cells in the standing orientation. Epifluorescence microscopy was used to image FCC cells post-labelling by the permeant DNA stain Syto9 (green channel). The scale bars in the small and large panels are 50 μm and 5 μm, respectively.

**Supplementary Table S1 – Statistical analysis from SDS-PAGE gels experiments on porin abundance in OMs.** All experiments were conducted for at least three biologically independent replicates. Technical replicates were averaged to produce replicate means that were subsequently used for analysis. Mean values were compared within and between groups using one-way ANOVA followed by Tukey’s post hoc for two-two comparisons. Differences were considered statistically significant if p<0.05 (True).

**Supplementary Table S2 – Statistical analysis from RT-qPCR experiments on porin expression in FCCs and SABs.** All experiments were conducted for at least three biologically independent replicates. Technical replicates were averaged to produce replicate means that were subsequently used for analysis. Mean values were compared within and between groups using one-way ANOVA followed by Tukey’s post hoc for two-two comparisons. Differences were considered statistically significant if p<0.05 (True).

**Supplementary Table S3 – Statistical analysis from DLS experiments on porin propensity to self-associate in DOTs.** All experiments were conducted for at least three biologically independent replicates. Technical replicates were averaged to produce replicate means that were subsequently used for analysis. Mean values were compared within and between groups using one-way ANOVA followed by Tukey’s post hoc for two-two comparisons. Differences were considered statistically significant if p<0.05 (True).

## Methods

### Strains

The *Providencia stuartii* ATCC 29914 strain was obtained from the Pasteur Institute (Paris, France). A disruption of the *omp-pst2* gene in *P. stuartii* ATCC 29914 (chloramphenicol-resistant)^22^ was performed to study the importance of Omp-Pst2 porin for adaptation to pathophysiological conditions. The *Escherichia coli* BL21 DE3 *ΔOmp8* strain^32,40^ – deleted of its principal porins OmpF, LamB, OmpA and OmpC – was used for the expression of *P. stuartii* porins for their purification prior to analyze their self-association capacity in DOTs by Dynamic Light Scattering (see detailed method below).

All chemicals were from Sigma-Aldrich, unless specified otherwise.

### Production of porins from P. stuartii in E. coli

One μl of plasmid pGOmp-Pst1 or pGOmp-Pst2 was transformed into 50 μL of *E. coli ΔOmp8* competent kanamycin-resistant cells by the heat-shock method^32,41^. Cells were recovered in 500 μL of SOC medium for 1 hour at 37°C. 20 μL of cell suspension was then transferred onto agar plates containing 100 and 25 ng.μL^-1^ ampicillin and kanamycin respectively, and incubated overnight at 37°C. A colony was introduced in 25 mL of Luria Bertani (LB) medium overnight, yielding a pre-culture that was further inoculated into 1 L of LB culture, selected by the addition of ampicillin and kanamycin at final concentrations of 100 μg/mL and 25 μg/mL, respectively. pGOmp-Pst1 and pGOmp-Pst2 plasmids contain a strong T7-phage promoter allowing sufficient porin expression without induction by IPTG. After the optical density of cultures at 600 nm reached a value of 1, cells were harvested by centrifugation at 4,500 rpm for 30 min and the pellets were immediately used for porins extraction.

### Porin extraction and solubilization

Pellet cells were resuspended in 20 mM phosphate buffer pH 7.4, supplemented with DNAse (few mg), 1 mM MgSO_4_ and an anti-protease cocktail (Complete, Roche). The cells were disrupted using a micro-fluidizer (Constant system LTD, UK) at 14,000 Psi. Membranes were separated from cytoplasm by centrifugation at 4,500 rpm for 45 min. The pellet containing the membranes was resuspended in 20 mM phosphate buffer pH 7.4 with 0,3% n-octyl-poly-oxyethylene (OPOE) detergent (Affymetrix, UK) and incubated at 20°C for 2 h under agitation to separate inner (IM) from outer membranes (OM). A 45-min ultracentrifugation at 35,000 rpm (Optima XE, Beckman Coulter) was then performed. The supernatant containing the OM was collected and the resulting pellet containing the IM and some remaining OM was resuspended and incubated at 20°C for 2 h in 20 mM phosphate buffer pH 7.4 with 3% OPOE. The last solubilization step was performed three times to maximize OM protein recovery. The supernatants containing the solubilized porins from the OM were stored at 4°C until purification.

### Porin purification

Solubilized porins were loaded onto an anionic-exchange column (Hitrap HQ, 5 mL, GE Healthcare Life Science, France) and subjected to detergent exchange with 0.1 M MES buffer pH 6.5 complemented with 0.1% lauryldimethylamine-oxide (LDAO) detergent (Affymetrix, UK) and 25 mM NaCl. The elimination of most lipopolysaccharides (LPS) was achieved by washing the column with 0.1 M MES pH 6.5 buffer complemented with 2% LDAO. After 4 h of slow washing (0.2 mL.min^-1^), the column was re-equilibrated with 0.1 M MES buffer pH 6.5 complemented with 0.1% LDAO and 25 mM NaCl. Proteins were then eluted by using of a NaCl gradient (0 to 1M). The fractions containing the pure protein were analyzed on SDS-PAGE to verify their purity. Those containing pure porin were pooled together and concentrated to 7 mg.mL^-1^ on a 100-kDa cutoff Amicon (Sigma-Aldrich, France) ultra-filtration unit. Purified porins were stored at 4°C until use.

### Liposome preparation

Large unilamellar liposomes (LUVs) were produced by standard film-hydration method^42^. Briefly, the liposome mixture was generated by adding 0.02% lissamine-rhodamine-sn-glycero-3-phosphoethanolamine (Rhod PE, Avanti Polar, Lipids Co., USA) to 4% L-a-phosphatidylcholine (egg PC, Avanti Polar, Lipids Co., USA) suspended in a chloroform solution. The lipids were dried under N2-flow to obtain a thin lipid film. Residual chloroform was eliminated by overnight vacuum. The lipid film was rehydrated with phosphate buffer (125 mM, pH 7.4) and vortexed continuously during 5 min to produce multilamellar vesicles. The vesicles were freeze-thawed (100K-310K) ten times to obtain LUVs. To finish, an extrusion with a 100-nm filter was performed to calibrate LUVs toward 70-nm diameter (mini-extruder with polycarbonate filters, Avanti Polar Lipids).

### Dynamic Light Scattering (DLS)

The hydrodynamic radius of LUVs was calculated after the incorporation of porins into their bilayer by using the DLS method (DynaPro Nanostar from Wyatt Technology). To this end, liposomes (0.125 mg.mL^-1^) and porins (ranged from 0.033 to 1.07 μM) were mixed and incubated at pH 7.4. To avoid a sudden decrease of detergent concentration in solution leading to a porin aggregation, we used biobeads (hydrophobic polystyrene) to decrease slightly the detergent below the critical micellar concentration (CMC). In the lipid bilayer, porins are oriented in natural way meaning oriented mostly with their intracellular turns facing the lumen of the LUVs and their extracellular loops facing the outside^43^. The incorporation of porins into liposomes generates the increase of hydrodynamic radius of liposomes continuing during 24 h at 4°C. The DLS measurement was then taken.

The most common habitat of *P. stuartii* in humans is the urinary tract. The aim of DLS study is to determine whether high concentrations of urea, ammonia (from 100 to 1,000 mM), bicarbonate, creatinine (from 10 to 100 mM) or pH variation (between 5 and 9) are compatible with the self-association of Omp-Pst1 and Omp-Pst2. For that purpose, stress conditions were applied before the incorporation of porins into LUVs. After 24 h incubation, biobeads were added and the hydrodynamic radii of proteoliposomes were determined another 24 h later.

### Transcriptomic study

*P. stuartii* forms FCC and SAB. Within both phenotypes, we aimed at monitoring the relative expression of Omp-Pst1 and Omp-Pst2 in *P. stuartii* ATCC 29914 and *P. stuartii*ΔOmp-Pst2 mutant when exposed to different concentrations of urea, ammonia and various pH.

#### Total RNA extraction

one colony was added to 30 mL of LB liquid medium and incubated at 37°C for 1 h under 150 rpm agitation. For each strain and each condition tested, three wells (triplicates) of a 6-well plate (CytoOne) were filled with 2 mL from the LB pre-culture. Plates were incubated at 37°C under 60 rpm agitation up to different OD at 600nm: 0.3, 0.5, 0.7, 0.9 or 1.5 to investigate porin expression at different growth phases. FCC in suspension were collected and briefly centrifuged to pellet the bacteria without damaging them. The pellet was resuspended in either 2 mL of fresh LB containing urea, ammonia (500 mM), bicarbonate (50 mM), or in 2 mL of LB medium pH-specific (between 5 and 9), and incubated at 37°C for 30 min under 60 rpm agitation. Concomitantly, adherent biofilms were washed with LB then 2 mL of fresh LB containing the stress conditions described above were added. The plate was incubated at 37°C for 30 min under 60 rpm agitation. The RNA extraction was performed using RNeasy Mini Kit (Qiagen) with some adjustments for the two different phenotypes in the primary steps. For FCC, 1 mL of the 2-mL culture was removed and replaced by 1 mL of RNAprotect Cell Reagent (Qiagen). After 5 min of incubation, the mixture was centrifuged at 3,200 g for 10 min and the pellet was resuspended in 110 μL of TE buffer (Tris-HCl 30 mM, EDTA 1 mM, pH 8.0) containing 15 mg/mL of lysozyme and 2 mg/mL of proteinase K (Qiagen), and incubated at RT for 10 min. For SAB, the 2 mL of LB medium were removed from well and 1 mL of RNAprotect Cell Reagent: LB (1: 1) was added. After 5 min incubation, the RNAprotect was removed and 410 uL of TE buffer containing 15 mg/mL of lysozyme and 0.5 mg/mL of proteinase K were added. The adherent biofilm was disrupted manually by scratching the well bottom and the solution was incubated at RT for 10 min. Total RNA from both phenotypes was extracted following manufacturer’s instructions. For both, quantity and quality of RNAs were assessed using a Nanodrop2000 (Thermo Fisher Scientific).

#### Reverse transcription quantitative PCR (RT-qPCR)

RNA extracted from FCC and SAB were immediately used for reverse transcription. One μg of total RNA was retro-transcribed into complementary DNA (cDNA) by using the QuantiTect Reverse Transcription Kit (Qiagen) following manufacturer’s instructions. cDNA was stored at −20°C until use and the remaining RNA were stored at −80°C.

The expressions of *omp-pst1* and *omp-pst2* were quantified by RT-qPCR. To perform an absolute quantification of transcript abundance of each porin in each condition tested, DNA fragments of known size and sequence were amplified and quantified from the genome of *P. stuartii* to be used as standards. To do so, *P. stuartii* genomic DNA was extracted using NucleoSpin Tissue kit (Macherey-Nagel) following manufacturer’s instructions. A 954 bp fragment of *omp-pst1* gene was amplified using the primers OmpPst1_F3qs (5’-GAAGATGGCGACGACTCACG-3’) and OmpPst1_R3qs (5’-GTAAACCAGACCCAGACCCAGAAC-3’), and a 956 bp fragment of *omp-pst2* gene was amplified using the primers OmpPst2_F4qs (5’-ATTATTCGCGGCGGGTGTTAC-3’) and OmpPst2_R4qs (5’-CAGCGGCCATATTCTTGTTGA-3’) using the Q5 High-Fidelity DNA Polymerase (New England BioLabs). The PCR program consisted in an initial step at 98°C for 5 min, then 35 cycles of 98°C for 30 sec (denaturation), 55°C for 30 sec (primers annealing) and 72°C for 2 min (elongation), ended by a final extension step at 72°C for 5 min. PCR amplicons were visualized on a 1 % agarose gel using SYBR Safe (Thermo Fisher Scientific). For each gene, the band corresponding to the amplicon was excised from the gel and purified using NucleoSpin Gel and PCR Clean-up kit (Macherey-Nagel) following manufacturer’s instructions. Quality and quantity were measured using a Nanodrop2000 (Thermo Fisher Scientific). Based on the sequence composition and length of each porin fragment amplified, the number of copies present in the standard was calculated and the standard curve was established using eight concentrations from 5.0 x10^1^ to 5.0 x10^8^ gene copies.

For the quantification of the copy number of each porin in each condition, a 77 bp fragment of *omp-pst1* gene was targeted using the primers OmpPst1_F1q (5’-CGCATTCGGTTATGTTGAT-3’) and OmpPst1_R1q (5’-CGCTTGACTTGTTGTTGT-3’), and a 82 bp fragment of *omp-pst2* gene using the primers OmpPst2_F2q (5’-CTTCGCTCTACAGTACCA-3’) and OmpPst2_R2q (5’-GCCATCACCATTGTTATCTAA-3’). The reaction consisted of 4 μL of ten fold-diluted samples or of the standards, 0.6 μL of each primer (final concentration of 10 μM each), 7 μL of SsoAdvanced Universal SYBR Green Supermix (Bio-Rad) and RNase-free water for a final reaction volume of 15 μL in each well. For each biological triplicate of each strain, two technical replicates were done. qPCR was run on a CFX Connect Real-Time PCR Detection System (Bio-Rad), and the qPCR program consisted in an initial step at 95°C for 3 min, then 39 cycles of 95°C for 10 sec (denaturation) and 60°C for 45 sec (primers annealing & elongation). A melt curve analysis was performed to assess the specificity of the amplification. The number of transcript copies of *omp-pst1* and *omp-pst2* was calculated using the standard curves for *P. stuartii* ATCC 29914 and *P. stuartii*ΔOmp-Pst2 by using Bio-Rad CFX Manager v.3.1 (Bio-Rad). Gene expression data was normalized using the total RNA content for each condition, which is the preferable normalization method when no housekeeping gene has been validated in the tested species, which is our case in *P. stuartii*^44^.

### Evaluation of porin abundance in the OM of P. stuartti cells

The abundance of porin proteins was quantified for *P. stuartii* ATCC 29914 and *P. stuartii*ΔOmp-Pst2 mutant under different conditions. One colony of each strain was inoculated in LB medium supplemented with 0, 100, 500 or 1,000 mM of urea, ammonia or with 0, 10, 25, 50 or 100 mM of bicarbonate, creatinine, or in pH-specific LB medium overnight. Ten microliters of bacteria from each condition was centrifuged for 30 s at 11,000 g, the pellet was resuspended in solution containing loading buffer 4X and 166 mM DDT, and heated to 95°C for 5 min. Cells denatured were then migrated on a SDS-PAGE gel for 90 min at 140 V. For porin quantification, the intensity of the corresponding band (between 35 and 40 kDa) was quantified and normalized by the total intensity of the profile using ImageJ software (v1.50i).

### Bacterial growth studies

First, we intended to understand the impact of the conditions from the human urinary tract on *P. stuartii* survival and ability to form FCC and SAB. We exposed *P. stuartii* to environmental cues such as urea, ammonia, bicarbonate, creatinine and pH variation. To this end, one colony of *P. stuartii* or *P. stuartii*ΔOmp-Pst2 (selected with 33 μg/mL of chloramphenicol) was inoculated in LB medium supplemented with 0, 100, 500 or 1,000 mM of urea, ammonia or with 0, 10, 25, 50 or 100 bicarbonate, creatinine, or in pH-specific LB medium for 1 h. Then, 150 μL of bacteria were disposed into a 96-well plate (Greiner) and incubated at 37°C under 100 rpm shaking overnight to promote biofilm formation on the well bottom and FCC in suspension. The bacterial growth was monitored by absorbance at 600 nm for 24 h (10 min interval between time points) using a Biotek Synergy H4 microplate reader (Winooski, VT, USA). Secondly, we investigated the effect of urea, ammonia, bicarbonate, creatinine and pH variation on FCC and SAB already established. We inoculated one colony of both *P. stuartii* strains in LB medium for 1 h. One hundred and fifty microliters of bacteria were then disposed into a 96-well plate and incubated overnight at 37°C under 100 rpm. The day after, FCC and SAB were exposed to 0, 100, 500 or 1,000 mM of urea, ammonia, or to 0, 10, 25, 50 or 100 bicarbonate, creatinine LB medium for 1 h, and the growth recovery was monitored over the day in terms of absorbance at 600 nm for 24h (10 minutes interval between time points) using a Biotek Synergy H4 microplate reader (Winooski, VT, USA).

### Imaging

To observe FCC and SAB, 7 μL of the planktonic bacteria were deposit between LB-Gelzan solid media (LB-Lennox solidified with 8 g/L Gelzan^TM^) and a glass cover-slide, and imaged immediately afterwards^45^. The wells were then washed with PBS three times to remove all floating bacteria and to image cells tightly attached to the well surface (= canonical biofilm). Bacteria were observed using an IX81 Olympus inverted microscope and samples were magnified thanks to a 100X objective (Plan APON100X, Olympus). The membranes were labeled using FM1-43X dye at 5 μg.ml^-1^ (479 nm excitation; emission: 598 nm emission), all bacteria (live and dead) were labeled using Syto® 9 Green at 5 μM (485 nm excitation; 498 nm emission) and only dead bacteria were labeled by propidium iodide solution at 20 μM (533 nm excitation; 617 nm emission). All fluorescent dyes were purchased from Thermo Scientific, USA.

### Statistical analysis

The *statsmodels* Python package was used for all analysis in this study^46^. We used the ‘Generalized Linear Model’ module. As we are dealing with positive, and continuous response variables, we chose the Gamma family, using an inverse law as link function. All factors and their correlation were included for a first fit, then insignificant correlations were removed until the most parsimonious model, containing all significant variables, was found – in less than 10 iterations on average, indicating a good convergence. The two porins were treated as independent throughout the process. Then, TukeyHSD test was used to perform post-hoc analyses on the most significant factors.

## Acknowledgements

We thank Martin Weik for continuous support, and Jean-Philippe Kleman, Françoise Lacroix, Aline Leroy and Anne-Marie Villard for their technical support during epifluorescence microscopy (VM4D plateform), DLS and RT-qPCR (RoBioMol plateform) experiments. Financial support by the Agence Nationale de la Recherche (ANR-15-CE18-0005 ClickBiofilm) and the GRAL labex (C7H-LXG11A20-DYNAMOP) is acknowledged, as well as support from the CEA, the CNRS and the Université Grenoble Alpes. This work used the platforms of the Grenoble Instruct Center (ISBG: UMS 3518 CNRS-CEA-UJF-EMBL) with support from FRISBI (ANR-10-INSB-05-02) and GRAL (ANR-10-LABX-49-01) within the Grenoble Partnership for Structural Biology (PSB).

## Author Contributions

J.L, G.T, M.E.K and J-P.C designed research and analyzed the data; J.L prepared samples and performed all experiments; G.T participated to RT-qPCR experiments; K.P. performed the statistical analyses of the data; M.E.K contributed preliminary data; J.L, G.T. and J.P.C. wrote the manuscript with input of all authors.

