## Supplementary Information for "Socialization of *Providencia stuartii* enables resistance to environmental insults"

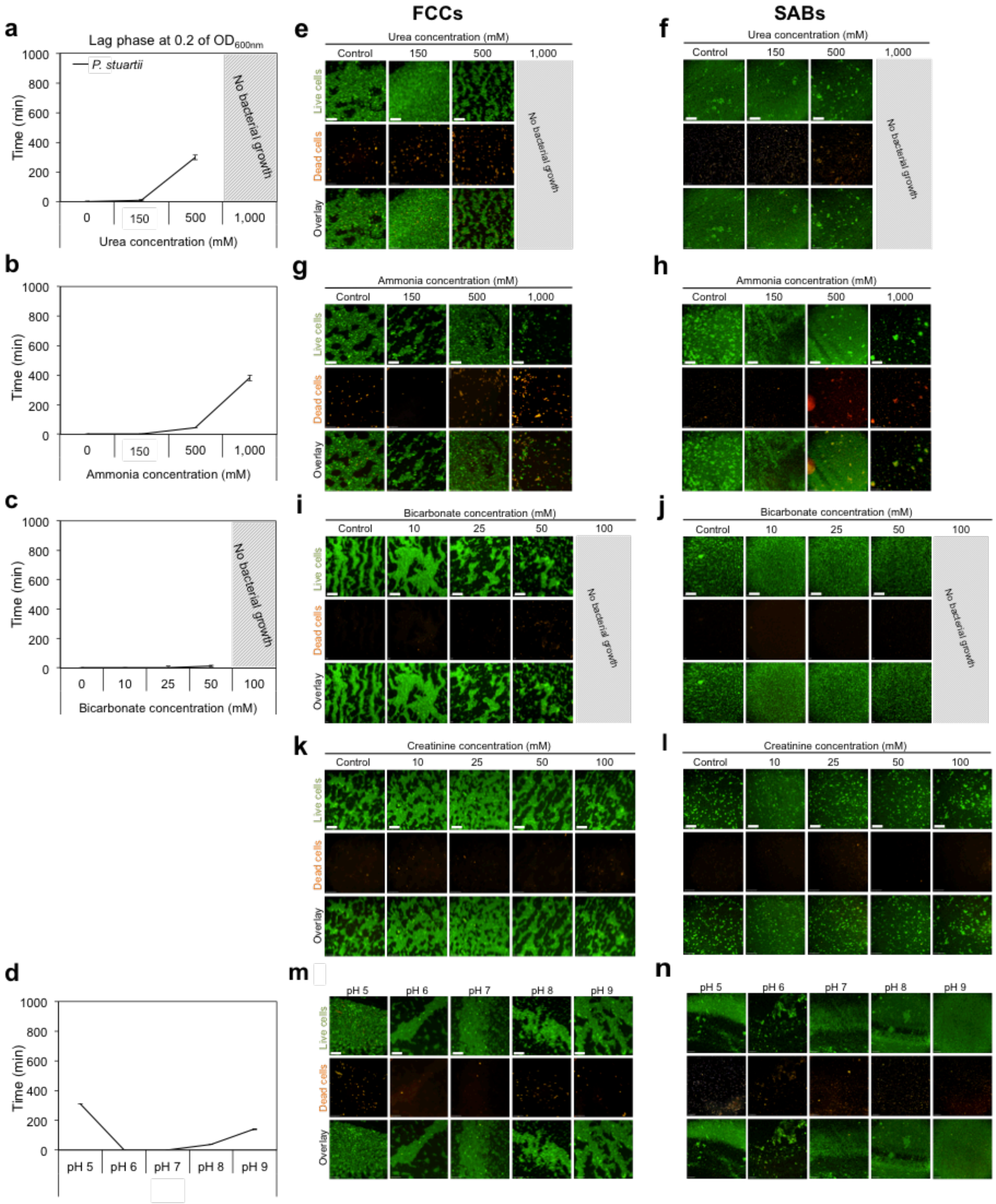


**Supplementary Figure S1. *P. stuartii* is highly resistant to catabolites present in the urinary tract. (a-d)** The overall impact of environmental cues on *P. stuartii* growth was monitored by determining, at increasing concentrations of the cues or pH, the lag time before reaching an optical density of 0.2 at 600 nm. Panels a, b, c, and d show results for urea, ammonia, bicarbonate and for different pH, respectively. All measurements were performed in triplicates; errors bars outline standard deviations from the average. Grey-hatched rectangles indicate concentrations at which bacteria were unable to grow. (**e-n**) Epifluorescence microscopy was used to monitor FCC (e,g,I,k,m) and SAB formation (f,h,j,l,n) in the presence of urea (e,f), ammonium (g,h), bicarbonate (i,j), creatinine (k,l) and at various pH (m,n). All bacteria are labelled by the permeant DNA stain Syto9 (green channel), but only dead cells are labelled by propidium iodide (red channel). For each condition, an overlay of the two channels is shown. Grey-hatched rectangles indicate concentrations at which bacteria were unable to grow. Scale bars correspond to 50 and 200 µm in FCC and SAB micrographs, respectively.

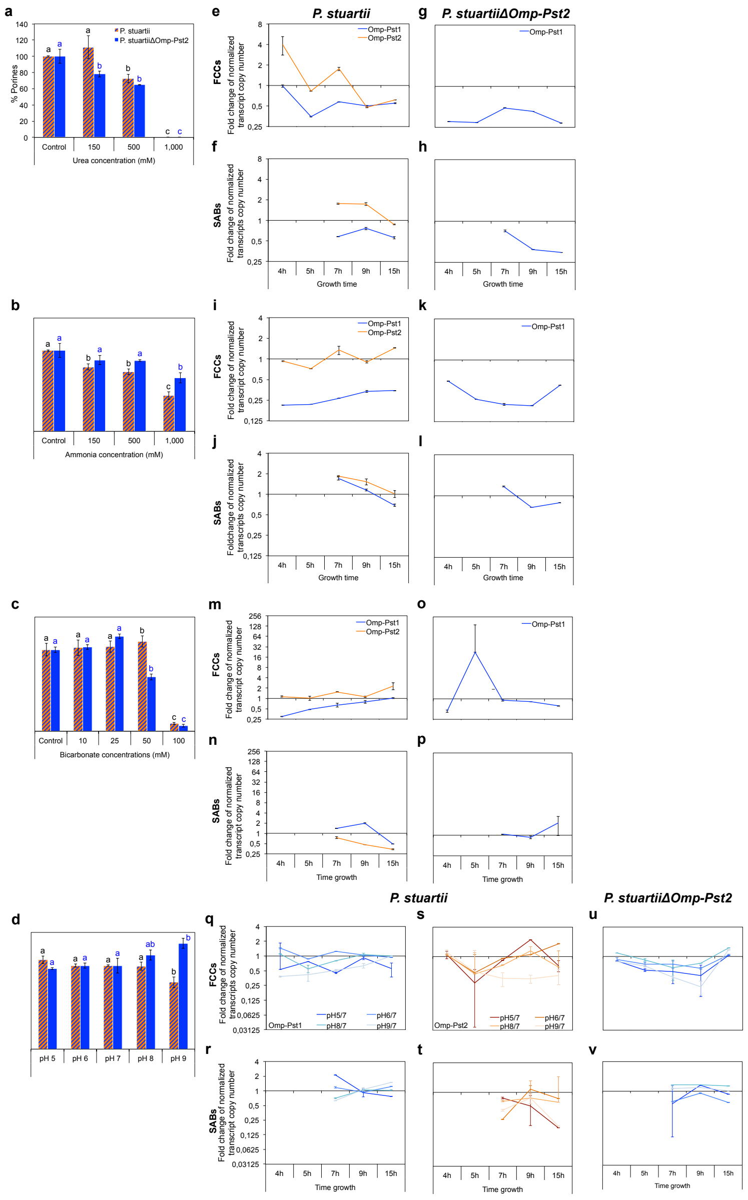


**Supplementary Figure S2. Regulation of porin expression in the presence of environmental cues is more pronounced in FCC than SAB, and sometimes opposed. (a-d)** Porin abundance in the OM of *P. stuartii* FCC and SAB cells grown in presence of increasing concentrations of urea (panel a), ammonia (b), bicarbonate (c) and to various pH (d) was evaluated by image processing of digitized SDS-PAGE gels using ImageJ. Plots show percent increase or decrease in porin abundance in the OM, after normalization of intensity counts from porin bands of exposed bacteria by those of unexposed bacteria. All measurements were performed in triplicates; errors bars outline standard deviations from the average. Different letters above the bars indicate significant differences ($p < 0.05$; ANOVA followed by post-hoc Tukey HSD test; see statistical indicators in Supplementary Table S1)**. (e-v)** Immediate changes in the expression of Omp-Pst1 (blue) and Omp-Pst2 (orange) in WT *P. stuartii* or *P. stuartii*ΔOmp-Pst2 FCCs and SABs cells were monitored by RT-qPCR following 30 min incubation with 500 mM of urea (panels e, f and g, h, respectively), 500 mM ammonia (i, j and k, l), to 50 mM bicarbonate (m, n and o, p), or to pH variation (q, r, s, t and u, v). For each growth time point, we report the ratio of normalized transcript copy numbers between cells exposed and not exposed (control) to the cues. All measurements were performed in triplicates; errors bars outline standard deviations from the average. Different letters above the bars indicate significant differences ($p < 0.05$; ANOVA followed by post-hoc Tukey HSD test; see statistical indicators in Supplementary Table S2)**.**

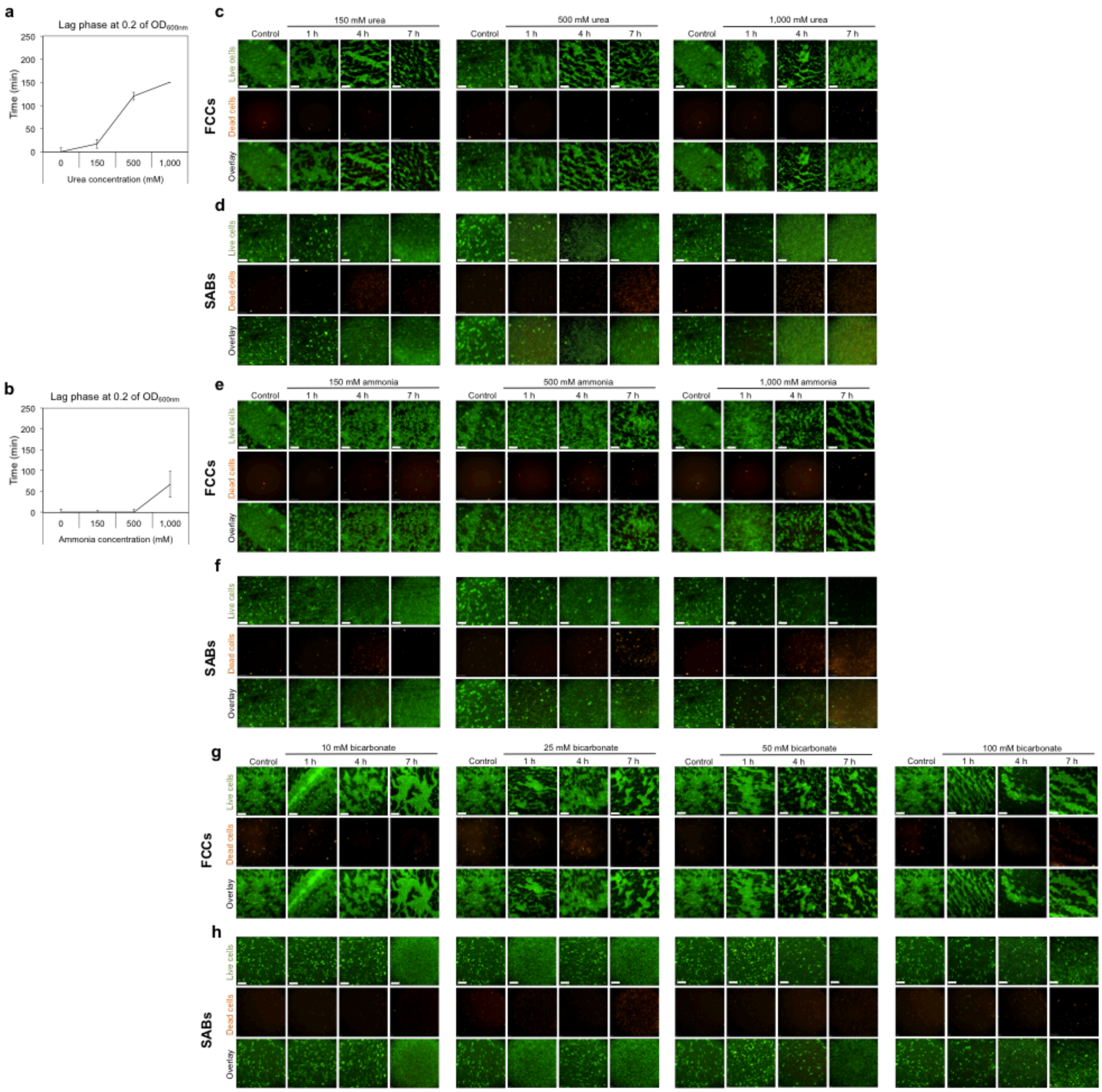


**Supplementary Figure S3. FCC and SAB cells are generally more resistant than their plantktonic counterparts. (a-b)** Recovery of preformed *P. stuartii* FCC and SAB cells after sudden exposure to urea (a) or ammonia (b) was monitored by determining, at increasing concentrations of these, the lag time before reaching an optical density of 0.2 at 600 nm. All measurements were performed in triplicates; errors bars outline standard deviations from the average. (**e-h**) Epifluorescence microscopy was used to monitor FCC (e,g,I,k,m) and SAB survival and recovery (f,h,j,l,n) in presence of increasing concentrations of urea (panels c and d, respectively), ammonia (e, f), and bicarbonate (g, h). All bacteria are labelled by the permeant DNA stain Syto9 (green channel), but only dead cells are labelled by propidium iodide (red channel). For each condition, an overlay of the two channels is shown. Grey-hatched rectangles indicate concentrations at which bacteria were unable to grow. Scale bars correspond to 50 and 200 µm in FCC and SAB micrographs, respectively. Note that preformed *P. stuartii* FCC and SAB generally resist better to the cues than their developing counterparts, as illustrated by survival of FCC and SAB at 1 M urea and 100 mM bicarbonate, where new cells do no grow (Supplementary Fig. 1). An exception is ammonium, which disrupts SAB formed in its absence although preformed FCC survive and new FCC and SAB can form (Supplementary Fig. 1).

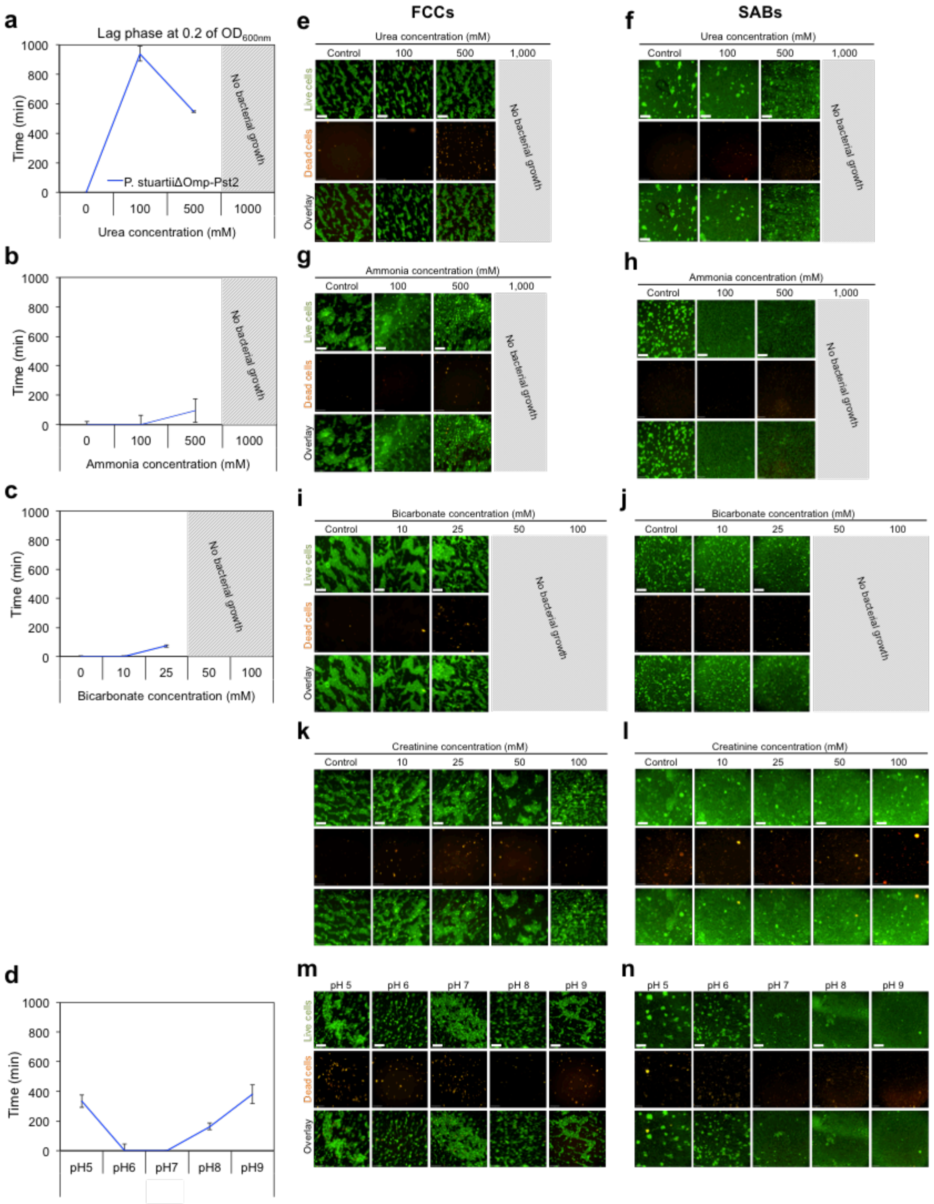

**Supplementary Figure S4. Omp-Pst2 benefits resistance of *P. stuartii* to its pathophysiological environment. (a-d)** The contribution of Omp-Pst2 in adapting to high concentration of urea (a), ammonia (b), bicarbonate (c) or to different pHs (d) was evaluated on the *P. stuartii*ΔOmp-Pst2 strain by determining, at increasing concentrations of the cues or pH, the lag time before reaching an optical density of 0.2 at 600 nm, and comparing results to the wild-type (Supplementary Fig. 1). All measurements were performed in triplicates; errors bars outline standard deviations from the average. Grey-hatched rectangles indicate concentrations at which bacteria were unable to grow. **(e-n)** Epifluorescence microscopy was used to monitor FCC (e,g,I,k,m) and SAB formation (f,h,j,l,n) by *P. stuartii*ΔOmp-Pst2 cells in the presence of urea (e,f), ammonium (g,h), bicarbonate (i,j), creatinine (k,l) and at various pH (m,n). All bacteria are labelled by the permeant DNA stain Syto9 (green channel), but only dead cells are labelled by propidium iodide (red channel). For each condition, an overlay of the two channels is shown. Grey-hatched rectangles indicate concentrations at which bacteria were unable to grow. Scale bars correspond to 50 and 200 µm in FCC and SAB micrographs, respectively.

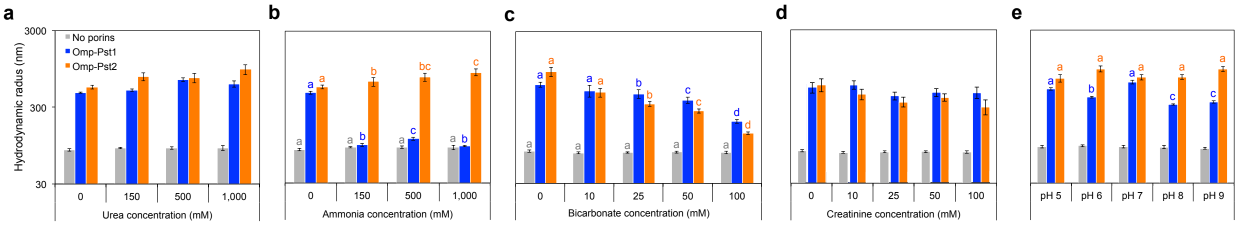


**Supplementary Figure S5. Omp-Pst1 and Omp-Pst2 self-association into DOT can be challenged by environmental cues.** LDAO-solubilized Omp-Pst1 (blue) and Omp-Pst2 (orange) were reconstituted in ~50-nm radius LUV, in presence of increasing concentrations of urea (a), ammonia (b), bicarbonate (c), creatinine (d) or at various pH (e), and after 24 h incubation with biobeads, the hydrodynamic radius of proteoliposomes was measured by DLS. The hydrodynamic radii of LUVs incubated at with the cues but without porin were also measured (grey). Different letters above the bars indicate significant differences ($p < 0.05$; ANOVA followed by post-hoc Tukey HSD test; see statistical indicators in Supplementary Table S3).

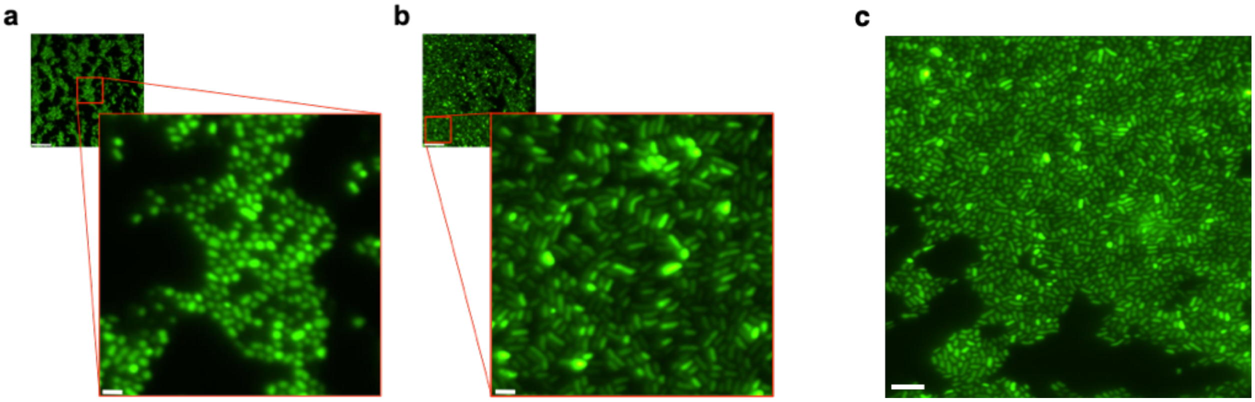


**Supplementary Figure S6. The orientation of *P. stuartii* cells differs in native FCC and in FCC formed in presence of environmental cues.** Close-up views of FCCs developed in the presence of urea (a) and ammonia (b), showing “standing” and “seated” orientations of cells, respectively. (c) Unexposed FCC feature cells in the standing orientation. Epifluorescence microscopy was used to image FCC cells post-labelling by the permeant DNA stain Syto9 (green channel). The scale bars in the small and large panels are 50 µm and 5 µm, respectively.

**Supplementary Table S1 – Statistical analysis from SDS-PAGE gels experiments on porin abundance in OMs.** All experiments were conducted for at least three biologically independent replicates. Technical replicates were averaged to produce replicate means that were subsequently used for analysis. Mean values were compared within and between groups using one-way ANOVA followed by Tukey's post hoc for two-two comparisons. Differences were considered statistically significant if p<0.05 (True).

#### GLM –SDS-PAGE Gels

##### All statistic parameters – *P. stuartii*:

```
                 Generalized Linear Model Regression Results
==============================================================================
Dep. Variable:                      Y   No. Observations:                   39
Model:                            GLM   Df Residuals:                       35
Model Family:                   Gamma   Df Model:                            3
Link Function:          inverse_power   Scale:                         0.20020
Method:                          IRLS   Log-Likelihood:                    inf
Date:                Thu, 14 Feb 2019   Deviance:                       212.99
Time:                        16:59:38   Pearson chi2:                     7.01
No. Iterations:                     8   Covariance Type:             nonrobust
=====================================================================================
                       coef    std err          z      P>|z|      [0.025      0.975]
-------------------------------------------------------------------------------------
Intercept            0.4455      0.395      1.129      0.259      -0.328       1.219
Urea              4.854e-05   1.21e-05      4.011      0.000    2.48e-05    7.23e-05
Ammonium          1.704e-05   6.75e-06      2.524      0.012    3.81e-06    3.03e-05
Urea:Ammonium     8.351e-16   7.63e-16      1.094      0.274   -6.61e-16    2.33e-15
pH                   0.0362      0.057      0.638      0.523      -0.075       0.147
Urea:pH              0.0003   8.47e-05      4.011      0.000       0.000       0.001
Ammonium:pH          0.0001   4.73e-05      2.524      0.012    2.67e-05       0.000
Urea:Ammonium:pH          0          0        nan        nan           0           0
=====================================================================================
```

##### Bicarbonate – *P. stuartii*:

```
                 Generalized Linear Model Regression Results
==============================================================================
Dep. Variable:                      Y   No. Observations:                   15
Model:                            GLM   Df Residuals:                       13
Model Family:                   Gamma   Df Model:                            1
Link Function:          inverse_power   Scale:                         0.29338
Method:                          IRLS   Log-Likelihood:                -136.43
Date:                Mon, 20 May 2019   Deviance:                       5.1540
Time:                        09:29:53   Pearson chi2:                     3.81
No. Iterations:                     7   Covariance Type:             nonrobust
=================================================================================
                    coef    std err          z      P>|z|      [0.025      0.975]
---------------------------------------------------------------------------------
Intercept         0.0001   3.41e-05      4.046      0.000    7.11e-05       0.000
Concentration  3.294e-06   1.26e-06      2.606      0.009    8.17e-07    5.77e-06
=================================================================================
```

##### All statistic parameters – *P. stuartii*ΔOmp-Pst2:

```
                 Generalized Linear Model Regression Results
==============================================================================
Dep. Variable:                 Values   No. Observations:                   39
Model:                            GLM   Df Residuals:                       35
Model Family:                   Gamma   Df Model:                            3
Link Function:          inverse_power   Scale:                         0.19051
Method:                          IRLS   Log-Likelihood:                    inf
Date:                Mon, 20 May 2019   Deviance:                       212.67
Time:                        09:37:42   Pearson chi2:                     6.67
No. Iterations:                     8   Covariance Type:             nonrobust
==============================================================================
                 coef    std err          z      P>|z|      [0.025      0.975]
------------------------------------------------------------------------------
Intercept      1.2797      0.498      2.572      0.010       0.305       2.255
Urea           0.0041      0.001      4.534      0.000       0.002       0.006
Ammonia        0.0005      0.000      1.670      0.095   -9.18e-05       0.001
pH            -0.0586      0.069     -0.855      0.392      -0.193       0.076
==============================================================================
```

##### Bicarbonate – *P. stuartii*ΔOmp-Pst2:

```
                 Generalized Linear Model Regression Results
==============================================================================
```

```
Dep. Variable:                      Y   No. Observations:                   15
Model:                            GLM   Df Residuals:                       13
Model Family:                   Gamma   Df Model:                            1
Link Function:          inverse_power   Scale:                         0.31456
Method:                          IRLS   Log-Likelihood:                -133.53
Date:                Mon, 20 May 2019   Deviance:                       7.3202
Time:                        09:30:46   Pearson chi2:                     4.09
No. Iterations:                     7   Covariance Type:             nonrobust
=================================================================================
                    coef    std err          z      P>|z|      [0.025      0.975]
---------------------------------------------------------------------------------
Intercept         0.0001   3.93e-05      3.569      0.000    6.33e-05       0.000
Concentration  7.247e-06   2.08e-06      3.491      0.000    3.18e-06    1.13e-05
=================================================================================
```

#### 2-2 comparison (Tukey) – *P. stuartii*

##### Urea condition:

```
Multiple Comparison of Means - Tukey HSD,FWER=0.05
=============================================
group1 group2 meandiff  lower   upper  reject
---------------------------------------------
  0     150    0.1542  -0.1792  0.4876 False
  0     500   -0.3759  -0.7093 -0.0425  True
  0     1000  -1.3651  -1.6985 -1.0317  True
 150    500   -0.5301  -0.8635 -0.1966  True
 150    1000  -1.5193  -1.8527 -1.1858  True
 500    1000  -0.9892  -1.3226 -0.6558  True
```

##### Ammonia condition:

```
Multiple Comparison of Means - Tukey HSD,FWER=0.05
=============================================
group1 group2 meandiff  lower   upper  reject
---------------------------------------------
  0     150   -0.2778  -0.4343 -0.1212  True
  0     500   -0.3618  -0.5183 -0.2052  True
  0     1000  -0.7592  -0.9157 -0.6026  True
 150    500    -0.084  -0.2405  0.0726 False
 150    1000  -0.4814  -0.6379 -0.3248  True
 500    1000  -0.3974  -0.5539 -0.2408  True
```

##### Bicarbonate condition:

```
  Multiple Comparison of Means - Tukey HSD,FWER=0.05
=====================================================
group1 group2  meandiff    lower      upper    reject
-----------------------------------------------------
  0      10    975.8383   211.0188  1740.6579   True
  0      25    756.3667   -8.4529   1521.1862  False
  0      50    780.3097   15.4901   1545.1292   True
  0     100   -4409.2103 -5174.0299 -3644.3908  True
  10     25   -219.4717  -984.2912   545.3479  False
  10     50   -195.5287  -960.3482   569.2909  False
  10    100   -5385.0487 -6149.8682 -4620.2291  True
  25     50     23.943   -740.8765   788.7625  False
  25    100   -5165.577  -5930.3965 -4400.7575  True
  50    100    -5189.52  -5954.3395 -4424.7005  True
```

##### pH condition:

```
Multiple Comparison of Means - Tukey HSD,FWER=0.05
=============================================
group1 group2 meandiff  lower   upper  reject
---------------------------------------------
  5      6    -0.0942  -0.2838  0.0954 False
  5      7    -0.0976  -0.2871  0.092  False
  5      8    -0.1051  -0.2947  0.0844 False
  5      9    -0.3684   -0.558 -0.1788  True
  6      7    -0.0034  -0.1929  0.1862 False
  6      8    -0.0109  -0.2005  0.1786 False
  6      9    -0.2742  -0.4638 -0.0846  True
  7      8    -0.0076  -0.1972  0.182  False
```

| | | | | | |
|---|---|---|---|---|---|
| 7 | 9 | -0.2708 | -0.4604 | -0.0813 | True |
| 8 | 9 | -0.2632 | -0.4528 | -0.0737 | True |

#### 2-2 comparison (Tukey) – *P. stuartii*ΔOmp-Pst2

##### Urea condition:

Multiple Comparison of Means - Tukey HSD,FWER=0.05

| group1 | group2 | meandiff | lower | upper | reject |
|---|---|---|---|---|---|
| 0 | 150 | -0.2351 | -0.4253 | -0.045 | True |
| 0 | 500 | -0.3753 | -0.5654 | -0.1851 | True |
| 0 | 1000 | -1.0683 | -1.2585 | -0.8782 | True |
| 150 | 500 | -0.1401 | -0.373 | 0.0928 | False |
| 150 | 1000 | -0.8332 | -1.0661 | -0.6003 | True |
| 500 | 1000 | -0.6931 | -0.926 | -0.4602 | True |

##### Ammonia condition:

Multiple Comparison of Means - Tukey HSD,FWER=0.05

| group1 | group2 | meandiff | lower | upper | reject |
|---|---|---|---|---|---|
| 0 | 150 | -0.1283 | -0.3409 | 0.0842 | False |
| 0 | 500 | -0.135 | -0.3475 | 0.0775 | False |
| 0 | 1000 | -0.3613 | -0.5738 | -0.1487 | True |
| 150 | 500 | -0.0066 | -0.2669 | 0.2536 | False |
| 150 | 1000 | -0.2329 | -0.4932 | 0.0274 | False |
| 500 | 1000 | -0.2263 | -0.4865 | 0.034 | False |

##### Bicarbonate condition:

Multiple Comparison of Means - Tukey HSD,FWER=0.05

| group1 | group2 | meandiff | lower | upper | reject |
|---|---|---|---|---|---|
| 0 | 10 | 346.731 | -289.3837 | 982.8457 | False |
| 0 | 25 | 154.9093 | -481.2053 | 791.024 | False |
| 0 | 50 | -2002.9833 | -2639.098 | -1366.8687 | True |
| 0 | 100 | -4633.1367 | -5269.2513 | -3997.022 | True |
| 10 | 25 | -191.8217 | -827.9363 | 444.293 | False |
| 10 | 50 | -2349.7143 | -2985.829 | -1713.5997 | True |
| 10 | 100 | -4979.8677 | -5615.9823 | -4343.753 | True |
| 25 | 50 | -2157.8927 | -2794.0073 | -1521.778 | True |
| 25 | 100 | -4788.046 | -5424.1607 | -4151.9313 | True |
| 50 | 100 | -2630.1533 | -3266.268 | -1994.0387 | True |

##### pH condition:

Multiple Comparison of Means - Tukey HSD,FWER=0.05

| group1 | group2 | meandiff | lower | upper | reject |
|---|---|---|---|---|---|
| 5 | 6 | 0.0374 | -0.2237 | 0.2985 | False |
| 5 | 7 | 0.0388 | -0.1744 | 0.252 | False |
| 5 | 8 | 0.1749 | -0.0862 | 0.436 | False |
| 5 | 9 | 0.3246 | 0.0635 | 0.5857 | True |
| 6 | 7 | 0.0014 | -0.2118 | 0.2146 | False |
| 6 | 8 | 0.1375 | -0.1236 | 0.3986 | False |
| 6 | 9 | 0.2872 | 0.0261 | 0.5483 | True |
| 7 | 8 | 0.1361 | -0.0771 | 0.3493 | False |
| 7 | 9 | 0.2858 | 0.0726 | 0.499 | True |
| 8 | 9 | 0.1497 | -0.1114 | 0.4108 | False |

**Supplementary Table S2 – Statistical analysis from RT-qPCR experiments on porin expression in FCCs and SABs.** All experiments were conducted for at least three biologically independent replicates. Technical replicates were averaged to produce replicate means that were subsequently used for analysis. Mean values were compared within and between groups using one-way ANOVA followed by Tukey's post hoc for two-two comparisons. Differences were considered statistically significant if $p<0.05$ (True).

### GLM – RTqPCR – *P. stuartii*

#### Parameter correlations: Urea - Omp-Pst1

```
                 Generalized Linear Model Regression Results
==============================================================================
Dep. Variable:                      Y   No. Observations:                   88
Model:                            GLM   Df Residuals:                       80
Model Family:                   Gamma   Df Model:                            7
Link Function:          inverse_power   Scale:                         0.22000
Method:                          IRLS   Log-Likelihood:                -1032.9
Date:                Tue, 21 May 2019   Deviance:                       19.302
Time:                        15:24:28   Pearson chi2:                     17.6
No. Iterations:                     7   Covariance Type:             nonrobust
======================================================================================================
                                         coef    std err          z      P>|z|      [0.025      0.975]
------------------------------------------------------------------------------------------------------
Intercept                           1.976e-05   4.34e-06      4.553      0.000    1.13e-05    2.83e-05
C(Phenotype)[T.Floating]           -5.015e-06   4.73e-06     -1.060      0.289   -1.43e-05    4.26e-06
Time                               -5.654e-07   3.72e-07     -1.518      0.129    -1.3e-06    1.64e-07
C(Phenotype)[T.Floating]:Time       1.242e-07   4.13e-07      0.301      0.763   -6.85e-07    9.33e-07
Urea                               -8.492e-09   3.04e-08     -0.279      0.780   -6.81e-08    5.11e-08
C(Phenotype)[T.Floating]:Urea       1.565e-08   3.27e-08      0.478      0.632   -4.85e-08    7.98e-08
Time:Urea                           3.323e-09   3.17e-09      1.049      0.294   -2.89e-09    9.53e-09
C(Phenotype)[T.Floating]:Time:Urea -1.773e-09   3.47e-09     -0.511      0.609   -8.57e-09    5.03e-09
======================================================================================================
```

#### Parameter correlations: Urea - Omp-Pst2

```
                 Generalized Linear Model Regression Results
==============================================================================
Dep. Variable:                      Y   No. Observations:                   88
Model:                            GLM   Df Residuals:                       80
Model Family:                   Gamma   Df Model:                            7
Link Function:          inverse_power   Scale:                         0.78223
Method:                          IRLS   Log-Likelihood:                -881.32
Date:                Tue, 21 May 2019   Deviance:                       56.367
Time:                        15:25:17   Pearson chi2:                     62.6
No. Iterations:                     8   Covariance Type:             nonrobust
======================================================================================================
                                         coef    std err          z      P>|z|      [0.025      0.975]
------------------------------------------------------------------------------------------------------
Intercept                              0.0001   4.72e-05      2.651      0.008    3.26e-05       0.000
C(Phenotype)[T.Floating]            8.957e-05   7.12e-05      1.257      0.209      -5e-05       0.000
Time                                -4.41e-06   3.91e-06     -1.127      0.260   -1.21e-05    3.26e-06
C(Phenotype)[T.Floating]:Time      -1.412e-06   6.45e-06     -0.219      0.827    -1.4e-05    1.12e-05
Urea                               -1.312e-07   1.32e-07     -0.995      0.320    -3.9e-07    1.27e-07
C(Phenotype)[T.Floating]:Urea      -9.149e-08   2.12e-07     -0.431      0.666   -5.08e-07    3.25e-07
Time:Urea                            6.08e-09   1.15e-08      0.530      0.596   -1.64e-08    2.86e-08
C(Phenotype)[T.Floating]:Time:Urea  1.064e-08   2.18e-08      0.488      0.625   -3.21e-08    5.33e-08
======================================================================================================
```

#### Parameter correlations: Ammonia - Omp-Pst1

```
                 Generalized Linear Model Regression Results
==============================================================================
Dep. Variable:                      Y   No. Observations:                   89
Model:                            GLM   Df Residuals:                       81
Model Family:                   Gamma   Df Model:                            7
Link Function:          inverse_power   Scale:                         0.18900
Method:                          IRLS   Log-Likelihood:                -1039.4
Date:                Tue, 21 May 2019   Deviance:                       15.403
Time:                        15:26:44   Pearson chi2:                     15.3
No. Iterations:                     7   Covariance Type:             nonrobust
==========================================================================================================
                                             coef    std err          z      P>|z|      [0.025      0.975]
----------------------------------------------------------------------------------------------------------
Intercept                               1.976e-05   4.02e-06      4.913      0.000    1.19e-05    2.76e-05
C(Phenotype)[T.Floating]               -5.015e-06   4.39e-06     -1.143      0.253   -1.36e-05    3.58e-06
Time                                   -5.654e-07   3.45e-07     -1.638      0.101   -1.24e-06    1.11e-07
C(Phenotype)[T.Floating]:Time           1.242e-07   3.82e-07      0.325      0.745   -6.25e-07    8.74e-07
```

```
Ammonium                                -2.697e-08   1.19e-08     -2.259      0.024   -5.04e-08   -3.57e-09
C(Phenotype)[T.Floating]:Ammonium        8.771e-08   2.01e-08      4.364      0.000    4.83e-08    1.27e-07
Time:Ammonium                             1.78e-09   1.11e-09      1.601      0.109   -3.99e-10    3.96e-09
C(Phenotype)[T.Floating]:Time:Ammonium -4.559e-09     1.8e-09     -2.530      0.011   -8.09e-09   -1.03e-09
============================================================================================================
```

#### Parameter correlations: Ammonia - Omp-Pst2

```
                 Generalized Linear Model Regression Results
==============================================================================
Dep. Variable:                      Y   No. Observations:                   89
Model:                            GLM   Df Residuals:                       81
Model Family:                   Gamma   Df Model:                            7
Link Function:          inverse_power   Scale:                         0.74485
Method:                          IRLS   Log-Likelihood:                -869.64
Date:                Tue, 21 May 2019   Deviance:                       51.910
Time:                        15:27:33   Pearson chi2:                     60.3
No. Iterations:                     8   Covariance Type:             nonrobust
============================================================================================================
                                             coef    std err          z      P>|z|      [0.025      0.975]
------------------------------------------------------------------------------------------------------------
Intercept                                  0.0001   4.61e-05      2.717      0.007    3.49e-05       0.000
C(Phenotype)[T.Floating]                8.957e-05   6.95e-05      1.289      0.198   -4.67e-05       0.000
Time                                    -4.41e-06   3.82e-06     -1.155      0.248   -1.19e-05    3.07e-06
C(Phenotype)[T.Floating]:Time          -1.412e-06   6.29e-06     -0.225      0.822   -1.37e-05    1.09e-05
Ammonium                                1.805e-07    2.8e-07      0.644      0.520   -3.69e-07     7.3e-07
C(Phenotype)[T.Floating]:Ammonium       4.068e-07   4.24e-07      0.961      0.337   -4.23e-07    1.24e-06
Time:Ammonium                          -5.772e-09    2.3e-08     -0.251      0.802   -5.08e-08    3.93e-08
C(Phenotype)[T.Floating]:Time:Ammonium -3.723e-08   3.37e-08     -1.106      0.269   -1.03e-07    2.87e-08
============================================================================================================
```

#### Parameter correlations: Bicarbonate - Omp-Pst1

```
                 Generalized Linear Model Regression Results
==============================================================================
Dep. Variable:                 Values   No. Observations:                   48
Model:                            GLM   Df Residuals:                       40
Model Family:                   Gamma   Df Model:                            7
Link Function:          inverse_power   Scale:                  0.28273516624
Method:                          IRLS   Log-Likelihood:                -618.86
Date:                Fri, 05 Jul 2019   Deviance:                       12.855
Time:                        13:51:13   Pearson chi2:                     11.3
No. Iterations:                     8
==============================================================================================================
                                                coef    std err          z      P>|z|      [95.0% Conf. Int.]
--------------------------------------------------------------------------------------------------------------
Intercept                                  1.779e-05    4.1e-06      4.341      0.000      9.76e-06  2.58e-05
C(Phenotypes)[T.Floating]                 -1.118e-05   4.31e-06     -2.591      0.010     -1.96e-05 -2.72e-06
Time                                      -1.128e-06   2.75e-07     -4.098      0.000     -1.67e-06 -5.88e-07
C(Phenotypes)[T.Floating]:Time              8.29e-07   2.97e-07      2.791      0.005      2.47e-07  1.41e-06
Cconditions                               -1.554e-08   9.83e-09     -1.581      0.114     -3.48e-08  3.73e-09
C(Phenotypes)[T.Floating]:Cconditions      3.568e-08   1.16e-08      3.063      0.002      1.29e-08  5.85e-08
Time:Cconditions                           1.162e-09   6.73e-10      1.728      0.084     -1.56e-10  2.48e-09
C(Phenotypes)[T.Floating]:Time:Cconditions -2.541e-09   8.11e-10     -3.133      0.002     -4.13e-09 -9.51e-10
==============================================================================================================
```

#### Parameter correlations: Bicarbonate - Omp-Pst2

```
                 Generalized Linear Model Regression Results
==============================================================================
Dep. Variable:                 Values   No. Observations:                   48
Model:                            GLM   Df Residuals:                       40
Model Family:                   Gamma   Df Model:                            7
Link Function:          inverse_power   Scale:                 0.350373625544
Method:                          IRLS   Log-Likelihood:                -516.90
Date:                Fri, 05 Jul 2019   Deviance:                       15.804
Time:                        14:02:51   Pearson chi2:                     14.0
No. Iterations:                     7
==============================================================================================================
                                                coef    std err          z      P>|z|      [95.0% Conf. Int.]
--------------------------------------------------------------------------------------------------------------
Intercept                                  2.502e-05   2.08e-05      1.204      0.229     -1.57e-05  6.58e-05
C(Phenotypes)[T.Floating]                 -3.345e-05   2.65e-05     -1.262      0.207     -8.54e-05  1.85e-05
Time                                       7.925e-07   2.01e-06      0.394      0.694     -3.15e-06  4.74e-06
C(Phenotypes)[T.Floating]:Time              7.57e-06   3.53e-06      2.143      0.032      6.48e-07  1.45e-05
Cconditions                               -8.522e-08   9.79e-08     -0.870      0.384     -2.77e-07  1.07e-07
C(Phenotypes)[T.Floating]:Cconditions      1.389e-07   1.06e-07      1.312      0.190     -6.87e-08  3.46e-07
Time:Cconditions                           1.602e-08    1.1e-08      1.452      0.146      -5.6e-09  3.76e-08
C(Phenotypes)[T.Floating]:Time:Cconditions -2.859e-08   1.29e-08     -2.221      0.026     -5.38e-08 -3.36e-09
==============================================================================================================
```

**Parameter correlations: pH - Omp-Pst1**

```
                 Generalized Linear Model Regression Results
==============================================================================
Dep. Variable:                      Y   No. Observations:                  156
Model:                            GLM   Df Residuals:                      148
Model Family:                   Gamma   Df Model:                            7
Link Function:          inverse_power   Scale:                         0.29369
Method:                          IRLS   Log-Likelihood:                -1843.2
Date:                Tue, 21 May 2019   Deviance:                       47.729
Time:                        15:29:23   Pearson chi2:                     43.5
No. Iterations:                     7   Covariance Type:             nonrobust
====================================================================================================
                                       coef    std err          z      P>|z|      [0.025      0.975]
----------------------------------------------------------------------------------------------------
Intercept                        -3.249e-05   2.38e-05     -1.365      0.172   -7.91e-05    1.42e-05
C(Phenotype)[T.Floating]          3.936e-05   2.59e-05      1.522      0.128   -1.13e-05    9.01e-05
Time                              3.894e-06   2.45e-06      1.593      0.111   -8.98e-07    8.69e-06
C(Phenotype)[T.Floating]:Time    -3.441e-06   2.61e-06     -1.317      0.188   -8.56e-06    1.68e-06
pH                                7.667e-06   3.44e-06      2.227      0.026     9.2e-07    1.44e-05
C(Phenotype)[T.Floating]:pH      -5.688e-06   3.74e-06     -1.523      0.128    -1.3e-05    1.63e-06
Time:pH                          -6.133e-07   3.43e-07     -1.786      0.074   -1.29e-06    5.97e-08
C(Phenotype)[T.Floating]:Time:pH   4.35e-07   3.67e-07      1.186      0.236   -2.84e-07    1.15e-06
====================================================================================================
```

**Parameter correlations: pH - Omp-Pst2**

```
                 Generalized Linear Model Regression Results
==============================================================================
Dep. Variable:                      Y   No. Observations:                  155
Model:                            GLM   Df Residuals:                      147
Model Family:                   Gamma   Df Model:                            7
Link Function:          inverse_power   Scale:                         0.90306
Method:                          IRLS   Log-Likelihood:                -1510.8
Date:                Tue, 21 May 2019   Deviance:                       112.64
Time:                        15:28:36   Pearson chi2:                     133.
No. Iterations:                     8   Covariance Type:             nonrobust
====================================================================================================
                                       coef    std err          z      P>|z|      [0.025      0.975]
----------------------------------------------------------------------------------------------------
Intercept                         6.154e-05      0.000      0.222      0.825      -0.000       0.001
C(Phenotype)[T.Floating]             0.0002      0.000      0.511      0.609      -0.001       0.001
Time                              2.285e-06   2.74e-05      0.083      0.934   -5.14e-05     5.6e-05
C(Phenotype)[T.Floating]:Time    -2.104e-05   3.64e-05     -0.578      0.563   -9.23e-05    5.03e-05
pH                                  1.1e-05   3.96e-05      0.278      0.781   -6.65e-05    8.85e-05
C(Phenotype)[T.Floating]:pH      -1.724e-05   5.17e-05     -0.334      0.739      -0.000     8.4e-05
Time:pH                          -7.931e-07   3.88e-06     -0.204      0.838    -8.4e-06    6.82e-06
C(Phenotype)[T.Floating]:Time:pH  3.284e-06   5.26e-06      0.624      0.532   -7.02e-06    1.36e-05
====================================================================================================
```

#### GLM – RTqPCR – *P. stuartii*ΔOmp-Pst2

**Parameter correlations: Urea**

```
                 Generalized Linear Model Regression Results
==============================================================================
Dep. Variable:                      Y   No. Observations:                   96
Model:                            GLM   Df Residuals:                       88
Model Family:                   Gamma   Df Model:                            7
Link Function:          inverse_power   Scale:                         0.20053
Method:                          IRLS   Log-Likelihood:                -1144.3
Date:                Tue, 21 May 2019   Deviance:                       19.798
Time:                        16:17:17   Pearson chi2:                     17.6
No. Iterations:                     8   Covariance Type:             nonrobust
======================================================================================================
                                         coef    std err          z      P>|z|      [0.025      0.975]
------------------------------------------------------------------------------------------------------
Intercept                           5.906e-06   2.35e-06      2.509      0.012    1.29e-06    1.05e-05
C(Phenotype)[T.Floating]             1.85e-06   2.72e-06      0.681      0.496   -3.48e-06    7.18e-06
Time                                 2.43e-07   2.34e-07      1.038      0.299   -2.16e-07    7.02e-07
C(Phenotype)[T.Floating]:Time      -7.038e-08   2.84e-07     -0.248      0.804   -6.26e-07    4.86e-07
Urea                                -5.05e-08   2.39e-08     -2.115      0.034   -9.73e-08    -3.7e-09
C(Phenotype)[T.Floating]:Urea       6.131e-08   2.77e-08      2.215      0.027    7.06e-09    1.16e-07
Time:Urea                           8.384e-09   2.96e-09      2.835      0.005    2.59e-09    1.42e-08
C(Phenotype)[T.Floating]:Time:Urea -5.163e-09   3.54e-09     -1.459      0.144   -1.21e-08    1.77e-09
======================================================================================================
```

#### Parameter correlations: Ammonia

```
                 Generalized Linear Model Regression Results
==============================================================================
Dep. Variable:                      Y   No. Observations:                   96
Model:                            GLM   Df Residuals:                       88
Model Family:                   Gamma   Df Model:                            7
Link Function:          inverse_power   Scale:                         0.15986
Method:                          IRLS   Log-Likelihood:                -1145.4
Date:                Tue, 21 May 2019   Deviance:                       15.809
Time:                        16:18:11   Pearson chi2:                     14.1
No. Iterations:                     7   Covariance Type:             nonrobust
============================================================================================================
                                             coef    std err          z      P>|z|      [0.025      0.975]
------------------------------------------------------------------------------------------------------------
Intercept                               5.906e-06    2.1e-06      2.810      0.005    1.79e-06       1e-05
C(Phenotype)[T.Floating]                 1.85e-06   2.43e-06      0.763      0.446   -2.91e-06    6.61e-06
Time                                     2.43e-07   2.09e-07      1.162      0.245   -1.67e-07    6.53e-07
C(Phenotype)[T.Floating]:Time          -7.038e-08   2.53e-07     -0.278      0.781   -5.67e-07    4.26e-07
Ammonium                               -7.175e-09   7.82e-09     -0.918      0.359   -2.25e-08    8.14e-09
C(Phenotype)[T.Floating]:Ammonium       5.693e-08   1.47e-08      3.863      0.000     2.8e-08    8.58e-08
Time:Ammonium                           5.944e-10   8.13e-10      0.731      0.465   -9.98e-10    2.19e-09
C(Phenotype)[T.Floating]:Time:Ammonium -2.649e-09   1.48e-09     -1.785      0.074   -5.56e-09    2.59e-10
============================================================================================================
```

#### Parameter correlations: Bicarbonate

```
                 Generalized Linear Model Regression Results
==============================================================================
Dep. Variable:                 Values   No. Observations:                   48
Model:                            GLM   Df Residuals:                       40
Model Family:                   Gamma   Df Model:                            7
Link Function:          inverse_power   Scale:                   1.13895772977
Method:                          IRLS   Log-Likelihood:                -612.73
Date:                Fri, 05 Jul 2019   Deviance:                       30.400
Time:                        15:15:31   Pearson chi2:                     45.6
No. Iterations:                     9
==================================================================================================
                                               coef    std err          z      P>|z|      [95.0% Conf. Int.]
--------------------------------------------------------------------------------------------------
Intercept                                 3.441e-05   1.61e-05      2.135      0.033      2.83e-06   6.6e-05
C(Phenotypes)[T.Floating]                -2.833e-06   1.91e-05     -0.148      0.882     -4.03e-05  3.46e-05
Time                                     -2.157e-06   1.09e-06     -1.988      0.047     -4.28e-06    -3e-08
C(Phenotypes)[T.Floating]:Time            1.842e-07   1.29e-06      0.143      0.886     -2.34e-06  2.71e-06
Cconditions                               8.339e-09   4.73e-08      0.176      0.860     -8.43e-08  1.01e-07
C(Phenotypes)[T.Floating]:Cconditions    -5.474e-08   5.21e-08     -1.051      0.293     -1.57e-07  4.74e-08
Time:Cconditions                         -7.075e-10   3.17e-09     -0.223      0.823     -6.92e-09  5.51e-09
C(Phenotypes)[T.Floating]:Time:Cconditions   4.06e-09   3.53e-09      1.149      0.250     -2.86e-09    1.1e-08
==================================================================================================
```

#### Parameter correlations: pH

```
                 Generalized Linear Model Regression Results
==============================================================================
Dep. Variable:                      Y   No. Observations:                  168
Model:                            GLM   Df Residuals:                      160
Model Family:                   Gamma   Df Model:                            7
Link Function:          inverse_power   Scale:                         0.21959
Method:                          IRLS   Log-Likelihood:                -2016.9
Date:                Tue, 21 May 2019   Deviance:                       34.902
Time:                        16:19:02   Pearson chi2:                     35.1
No. Iterations:                     7   Covariance Type:             nonrobust
======================================================================================================
                                       coef    std err          z      P>|z|      [0.025      0.975]
------------------------------------------------------------------------------------------------------
Intercept                         1.953e-05   1.05e-05      1.856      0.063    -1.1e-06    4.02e-05
C(Phenotype)[T.Floating]         -9.239e-06   1.27e-05     -0.725      0.468   -3.42e-05    1.57e-05
Time                             -2.693e-07   1.01e-06     -0.266      0.790   -2.25e-06    1.71e-06
C(Phenotype)[T.Floating]:Time      5.56e-07   1.27e-06      0.436      0.663   -1.94e-06    3.05e-06
pH                               -1.726e-06   1.41e-06     -1.222      0.222   -4.49e-06    1.04e-06
C(Phenotype)[T.Floating]:pH       2.052e-06   1.73e-06      1.183      0.237   -1.35e-06    5.45e-06
Time:pH                           6.412e-08   1.37e-07      0.469      0.639   -2.04e-07    3.32e-07
C(Phenotype)[T.Floating]:Time:pH -1.241e-07   1.74e-07     -0.712      0.476   -4.66e-07    2.17e-07
======================================================================================================
```

#### 2-2 comparison (Tukey) - Urea condition – *P. stuartii*

#### Omp-Pst1 porin – FCCs

```
        Multiple Comparison of Means - Tukey HSD,FWER=0.05
```

| | group1 | group2 | meandiff | lower | upper | reject |
|---|---|---|---|---|---|---|
| 4h | 0 | 500 | 16574.1498 | -30452.3034 | 63600.6031 | False |
| 5h | 0 | 500 | -38902.0202 | -69962.3523 | -7841.6881 | True |
| 7h | 0 | 500 | -71189.8961 | -123737.1743 | -18642.618 | True |
| 9h | 0 | 500 | -63057.0912 | -127634.1388 | 1519.9565 | True |
| 15h | 0 | 500 | -63286.2712 | -114749.5456 | -11822.9968 | True |

#### Omp-Pst2 porin – FCCs

Multiple Comparison of Means - Tukey HSD,FWER=0.05

| | group1 | group2 | meandiff | lower | upper | reject |
|---|---|---|---|---|---|---|
| 4h | 0 | 500 | 6261.4946 | 1996.197 | 10526.7923 | True |
| 5h | 0 | 500 | -2134.6555 | - 12996.94 | 8727.629 | False |
| 7h | 0 | 500 | 7521.8879 | 1598.1456 | 13445.6302 | True |
| 9h | 0 | 500 | -657.2522 | -10717.0073 | 9402.5029 | False |
| 15h | 0 | 500 | -539.4299 | -9813.6084 | 8734.7485 | False |

#### 2-2 comparison (Tukey) - Urea condition – *P. stuartii* – According Time

#### Omp-Pst1 porin – FCCs

Multiple Comparison of Means - Tukey HSD,FWER=0.05

| group1 | group2 | meandiff | lower | upper | reject |
|---|---|---|---|---|---|
| 4 | 5 | -69510.7473 | -90915.2466 | -48106.248 | True |
| 4 | 7 | -30288.1114 | -51692.6107 | -8883.6121 | True |
| 4 | 9 | -47009.2685 | -68413.7677 | -25604.7692 | True |
| 4 | 15 | -35263.6879 | -56668.1872 | -13859.1886 | True |
| 5 | 7 | 39222.6359 | 17818.1367 | 60627.1352 | True |
| 5 | 9 | 22501.4789 | 1096.9796 | 43905.9781 | True |
| 5 | 15 | 34247.0594 | 12842.5601 | 55651.5587 | True |
| 7 | 9 | -16721.1571 | -38125.6564 | 4683.3422 | False |
| 7 | 15 | -4975.5765 | -26380.0758 | 16428.9228 | False |
| 9 | 15 | 11745.5806 | -9658.9187 | 33150.0798 | False |

#### 2-2 comparison (Tukey) – Ammonia condition – *P. stuartii*

#### Omp-Pst1 porin – FCCs

Multiple Comparison of Means - Tukey HSD,FWER=0.05

| | group1 | group2 | meandiff | lower | upper | reject |
|---|---|---|---|---|---|---|
| 4H | 0 | 500 | -48214.2397 | -94654.66 | -1773.8194 | True |
| 5H | 0 | 500 | -35184.7269 | -66101.6521 | -4267.8018 | True |
| 7H | 0 | 500 | -80993.2267 | -134113.1314 | -27873.3221 | True |
| 9H | 0 | 500 | -54354.4792 | -118532.4498 | 9823.4914 | True |
| 15H | 0 | 500 | -63325.4663 | -115029.2888 | -11621.6438 | True |

#### Omp-Pst1 porin – SABs

Multiple Comparison of Means - Tukey HSD,FWER=0.05

| | group1 | group2 | meandiff | lower | upper | reject |
|---|---|---|---|---|---|---|
| 7H | 0 | 500 | 51763.2687 | 13226.7333 | 90299.804 | True |
| 9H | 0 | 500 | 44588.3453 | -2241.4323 | 91418.123 | False |
| 15H | 0 | 500 | 122.8988 | -69268.6729 | 69514.4706 | False |

#### 2-2 comparison (Tukey) – Bicarbonate condition – *P. stuartii*

#### Omp-Pst1 porin – FCCs

Multiple Comparison of Means - Tukey HSD,FWER=0.05

| | group1 | group2 | meandiff | lower | upper | reject |
|---|---|---|---|---|---|---|
| 4h | 0 | 50 | -252343.4867 | -407839.5819 | -96847.3915 | True |
| 5h | 0 | 50 | -54122.7379 | -107260.2379 | -985.2379 | True |
| 7h | 0 | 50 | -53049.4907 | -161346.8928 | 55247.9113 | False |
| 9h | 0 | 50 | -37971.1624 | -256101.1496 | 180158.8249 | False |
| 15h | 0 | 50 | 5929.5254 | -256043.3444 | 267902.3952 | False |

#### Omp-Pst2 porin – FCCs

Multiple Comparison of Means - Tukey HSD,FWER=0.05

| | group1 | group2 | meandiff | lower | upper | reject |
|---|---|---|---|---|---|---|
| 4h | 0 | 50 | 6895.8395 | -59887.8827 | 73679.5616 | False |
| 5h | 0 | 50 | 345.2546 | -17834.8625 | 18525.3718 | False |
| 7h | 0 | 50 | 11245.2606 | -8163.8192 | 30654.3404 | False |
| 9h | 0 | 50 | 877.444 | -9994.3841 | 11749.2721 | False |
| 15h | 0 | 50 | 16834.8001 | 595.501 | 33074.0991 | True |

#### 2-2 comparison (Tukey) – pH condition – *P. stuartii*

#### Omp-Pst1 porin – FCCs

### 4h:

Multiple Comparison of Means - Tukey HSD,FWER=0.05

| group1 | group2 | meandiff | lower | upper | reject |
|---|---|---|---|---|---|
| 5 | 6 | 30203.4498 | -33493.5242 | 93900.4238 | False |
| 5 | 7 | 49864.7744 | -2949.9663 | 102679.5152 | False |
| 5 | 8 | 18929.9289 | -44767.0451 | 82626.9029 | False |
| 5 | 9 | -5065.2398 | -68762.2138 | 58631.7342 | False |
| 6 | 7 | 19661.3246 | -33153.4161 | 72476.0654 | False |
| 6 | 8 | -11273.5209 | -74970.4949 | 52423.4531 | False |
| 6 | 9 | -35268.6896 | -98965.6636 | 28428.2843 | False |
| 7 | 8 | -30934.8455 | -83749.5863 | 21879.8952 | False |
| 7 | 9 | -54930.0143 | -107744.755 | -2115.2735 | True |
| 8 | 9 | -23995.1687 | -87692.1427 | 39701.8052 | False |

### 5h:

Multiple Comparison of Means - Tukey HSD,FWER=0.05

| group1 | group2 | meandiff | lower | upper | reject |
|---|---|---|---|---|---|
| 5 | 6 | 4324.7478 | -35825.5557 | 44475.0512 | False |
| 5 | 7 | 19422.9899 | -13867.883 | 52713.8628 | False |
| 5 | 8 | -9858.4211 | -50008.7245 | 30291.8823 | False |
| 5 | 9 | -15480.8529 | -55631.1564 | 24669.4505 | False |
| 6 | 7 | 15098.2422 | -18192.6308 | 48389.1151 | False |
| 6 | 8 | -14183.1688 | -54333.4722 | 25967.1346 | False |
| 6 | 9 | -19805.6007 | -59955.9041 | 20344.7027 | False |
| 7 | 8 | -29281.411 | -62572.2839 | 4009.4619 | False |
| 7 | 9 | -34903.8429 | -68194.7158 | -1612.9699 | True |
| 8 | 9 | -5622.4319 | -45772.7353 | 34527.8716 | False |

### 7h:

Multiple Comparison of Means - Tukey HSD,FWER=0.05

| group1 | group2 | meandiff | lower | upper | reject |
|---|---|---|---|---|---|
| 5 | 6 | 83222.8583 | 9312.0111 | 157133.7054 | True |
| 5 | 7 | 77210.3097 | 15926.6727 | 138493.9467 | True |
| 5 | 8 | 37275.7864 | -36635.0608 | 111186.6335 | False |
| 5 | 9 | 6370.9945 | -67539.8527 | 80281.8416 | False |
| 6 | 7 | -6012.5486 | -67296.1855 | 55271.0884 | False |
| 6 | 8 | -45947.0719 | -119857.919 | 27963.7752 | False |
| 6 | 9 | -76851.8638 | -150762.7109 | -2941.0167 | True |
| 7 | 8 | -39934.5233 | -101218.1603 | 21349.1136 | False |
| 7 | 9 | -70839.3152 | -132122.9522 | -9555.6783 | True |
| 8 | 9 | -30904.7919 | -104815.639 | 43006.0552 | False |

#### Omp-Pst1 porin – SABs

### 7h:

Multiple Comparison of Means - Tukey HSD,FWER=0.05

| group1 | group2 | meandiff | lower | upper | reject |
|---|---|---|---|---|---|
| 5 | 6 | -26759.8469 | -78909.5474 | 25389.8536 | False |
| 5 | 7 | -33941.5292 | -76521.5814 | 8638.5229 | False |
| 5 | 8 | -66739.4717 | -118889.1722 | -14589.7711 | True |
| 5 | 9 | -70736.6287 | -122886.3292 | -18586.9282 | True |
| 6 | 7 | -7181.6824 | -49761.7345 | 35398.3698 | False |
| 6 | 8 | -39979.6248 | -92129.3253 | 12170.0757 | False |
| 6 | 9 | -43976.7818 | -96126.4823 | 8172.9187 | False |
| 7 | 8 | -32797.9424 | -75377.9946 | 9782.1098 | False |
| 7 | 9 | -36795.0995 | -79375.1516 | 5784.9527 | False |
| 8 | 9 | -3997.157 | -56146.8576 | 48152.5435 | False |

#### 2-2 comparison (Tukey) - Urea condition – *P. stuartii*ΔOmp-Pst2

##### Omp-Pst1 porin – FCCs

Multiple Comparison of Means - Tukey HSD,FWER=0.05

| | group1 | group2 | meandiff | lower | upper | reject |
|---|---|---|---|---|---|---|
| 4h | 0 | 500 | -48814.8603 | -79244.6239 | -18385.0967 | True |
| 5h | 0 | 500 | -79571.3733 | -112223.3454 | -46919.4013 | True |
| 7h | 0 | 500 | -84041.5601 | -146780.2218 | -21302.8985 | True |
| 9h | 0 | 500 | -80294.8613 | -120605.1033 | -39984.6194 | True |
| 15h | 0 | 500 | -66788.6326 | -124584.7873 | -8992.4779 | True |

##### Omp-Pst1 porin – SABs

Multiple Comparison of Means - Tukey HSD,FWER=0.05

| | group1 | group2 | meandiff | lower | upper | reject |
|---|---|---|---|---|---|---|
| 7h | 0 | 500 | -51532.3838 | -131390.0291 | 28325.2614 | False |
| 9h | 0 | 500 | -67561.3871 | -143722.56 | 8599.7859 | False |
| 15h | 0 | 500 | -85432.9544 | -138213.5268 | -32652.382 | True |

#### 2-2 comparison (Tukey) – Ammonia condition – *P. stuartii*ΔOmp-Pst2

##### Omp-Pst1 porin – FCC

Multiple Comparison of Means - Tukey HSD,FWER=0.05

| | group1 | group2 | meandiff | lower | upper | reject |
|---|---|---|---|---|---|---|
| 4h | 0 | 500 | -40754.5817 | -70288.9453 | -11220.2181 | True |
| 5h | 0 | 500 | -79073.6582 | -112314.5001 | -45832.8162 | True |
| 7h | 0 | 500 | -104908.034 | -169132.0224 | -40684.0456 | True |
| 9h | 0 | 500 | -108527.7472 | -148262.142 | -68793.3525 | True |
| 15h | 0 | 500 | -21559.4458 | -79543.416 | 36424.5245 | True |

#### 2-2 comparison (Tukey) – pH condition – *P. stuartii*ΔOmp-Pst2

##### Omp-Pst1 porin – FCC

### 5h:

Multiple Comparison of Means - Tukey HSD,FWER=0.05

| group1 | group2 | meandiff | lower | upper | reject |
|---|---|---|---|---|---|
| 5 | 6 | 17463.0878 | -29975.2601 | 64901.4357 | False |
| 5 | 7 | 61544.038 | 22810.7891 | 100277.2868 | True |
| 5 | 8 | 31719.0228 | -15719.3251 | 79157.3706 | False |
| 5 | 9 | 11726.8916 | -35711.4563 | 59165.2394 | False |
| 6 | 7 | 44080.9502 | 5347.7013 | 82814.199 | True |
| 6 | 8 | 14255.935 | -33182.4129 | 61694.2828 | False |
| 6 | 9 | -5736.1962 | -53174.5441 | 41702.1516 | False |
| 7 | 8 | -29825.0152 | -68558.2641 | 8908.2336 | False |

```
  7      9    -49817.1464 -88550.3953 -11083.8976  True
  8      9    -19992.1312 -67430.4791  27446.2167 False
```

## 7h:

```
    Multiple Comparison of Means - Tukey HSD,FWER=0.05
=========================================================
group1 group2   meandiff     lower        upper    reject
---------------------------------------------------------
  5      6     25669.2245 -61218.4629  112556.9119 False
  5      7     82700.2695  11756.7698  153643.7692  True
  5      8     12770.8041 -74116.8833   99658.4916 False
  5      9    -14547.9893 -101435.6767  72339.6981 False
  6      7     57031.045  -13912.4547  127974.5447 False
  6      8    -12898.4204 -99786.1078   73989.267  False
  6      9    -40217.2138 -127104.9013  46670.4736 False
  7      8    -69929.4654 -140872.9651  1014.0343  False
  7      9    -97248.2588 -168191.7585 -26304.7591  True
  8      9    -27318.7935 -114206.4809  59568.894  False
```

## 9h:

```
    Multiple Comparison of Means - Tukey HSD,FWER=0.05
==========================================================
group1 group2   meandiff      lower        upper    reject
----------------------------------------------------------
  5      6     21948.695   -38626.9602   82524.3501 False
  5      7     84516.5196   35056.7043  133976.3349  True
  5      8     43319.6633  -17255.9918  103895.3184 False
  5      9    -22639.6063  -83215.2615   37936.0488 False
  6      7     62567.8246   13108.0093  112027.6399  True
  6      8     21370.9683  -39204.6868   81946.6235 False
  6      9    -44588.3013  -105163.9564  15987.3538 False
  7      8    -41196.8563  -90656.6716    8262.959  False
  7      9    -107156.1259 -156615.9412 -57696.3107  True
  8      9    -65959.2696  -126534.9248  -5383.6145  True
```

**Supplementary Table S3 – Statistical analysis from DLS experiments on porin propensity to self-associate in DOTs.** All experiments were conducted for at least three biologically independent replicates. Technical replicates were averaged to produce replicate means that were subsequently used for analysis. Mean values were compared within and between groups using one-way ANOVA followed by Tukey's post hoc for two-two comparisons. Differences were considered statistically significant if $p<0.05$ (True).

#### GLM - DLS

##### Omp-Pst1 porin – All statistic parameters:

```
                 Generalized Linear Model Regression Results
==============================================================================
Dep. Variable:                      Y   No. Observations:                  389
Model:                            GLM   Df Residuals:                      381
Model Family:                   Gamma   Df Model:                            7
Link Function:          inverse_power   Scale:                        0.097908
Method:                          IRLS   Log-Likelihood:                -1972.8
Date:                Mon, 18 Mar 2019   Deviance:                       33.696
Time:                        14:17:24   Pearson chi2:                     37.3
No. Iterations:                     7   Covariance Type:             nonrobust
===================================================================================
                      coef    std err          z      P>|z|      [0.025      0.975]
-----------------------------------------------------------------------------------
Intercept           0.0137      0.002      8.950      0.000       0.011       0.017
Cporin             -0.0130      0.002     -7.730      0.000      -0.016      -0.010
Urea               2.3e-07   6.93e-07      0.332      0.740   -1.13e-06    1.59e-06
Cporin:Urea      7.344e-07   8.34e-07      0.881      0.378      -9e-07    2.37e-06
Ammonium         3.751e-06   9.29e-07      4.036      0.000    1.93e-06    5.57e-06
Cporin:Ammonium   4.42e-06   1.36e-06      3.246      0.001    1.75e-06    7.09e-06
pH                 -0.0007      0.000     -3.151      0.002      -0.001      -0.000
Cporin:pH           0.0012      0.000      5.136      0.000       0.001       0.002
===================================================================================
```

##### Omp-Pst2 porin – All statistic parameters:

```
                 Generalized Linear Model Regression Results
==============================================================================
Dep. Variable:                      Y   No. Observations:                  386
Model:                            GLM   Df Residuals:                      378
Model Family:                   Gamma   Df Model:                            7
Link Function:          inverse_power   Scale:                         0.36485
Method:                          IRLS   Log-Likelihood:                -2501.6
Date:                Mon, 18 Mar 2019   Deviance:                       123.99
Time:                        14:19:20   Pearson chi2:                     138.
No. Iterations:                     8   Covariance Type:             nonrobust
===================================================================================
                      coef    std err          z      P>|z|      [0.025      0.975]
-----------------------------------------------------------------------------------
Intercept           0.0066      0.001      4.551      0.000       0.004       0.010
Cporin             -0.0046      0.001     -3.105      0.002      -0.008      -0.002
Urea             4.958e-07   6.87e-07      0.722      0.470   -8.51e-07    1.84e-06
Cporin:Urea     -6.935e-07   6.79e-07     -1.022      0.307   -2.02e-06    6.36e-07
Ammonium         1.284e-05   1.92e-06      6.681      0.000    9.07e-06    1.66e-05
Cporin:Ammonium -1.208e-05   1.82e-06     -6.651      0.000   -1.56e-05   -8.52e-06
pH                 -0.0003      0.000     -1.475      0.140      -0.001    9.77e-05
Cporin:pH           0.0002      0.000      1.107      0.268      -0.000       0.001
===================================================================================
```

##### Omp-Pst1 porin – Bicarbonate – All statistic parameters:

```
                 Generalized Linear Model Regression Results
==============================================================================
Dep. Variable:                      Y   No. Observations:                   96
Model:                            GLM   Df Residuals:                       92
Model Family:                   Gamma   Df Model:                            3
Link Function:          inverse_power   Scale:                       0.0066722
Method:                          IRLS   Log-Likelihood:                -380.86
Date:                Sat, 20 Apr 2019   Deviance:                      0.62057
Time:                        11:22:54   Pearson chi2:                    0.614
No. Iterations:                     7   Covariance Type:             nonrobust
=========================================================================================
                            coef    std err          z      P>|z|      [0.025      0.975]
-----------------------------------------------------------------------------------------
Intercept                 0.0129      0.000     60.130      0.000       0.012       0.013
Bicarbonate            2.458e-06   4.09e-06      0.601      0.548   -5.56e-06    1.05e-05
Concentration            -0.0109      0.000    -50.047      0.000      -0.011      -0.010
```

```
Bicarbonate:Concentration  1.985e-05   4.23e-06      4.691      0.000    1.16e-05    2.81e-05
=============================================================================================
```

#### Omp-Pst1 porin – Bicarbonate:

```
                 Generalized Linear Model Regression Results                  
==============================================================================
Dep. Variable:                      Y   No. Observations:                   96
Model:                            GLM   Df Residuals:                       94
Model Family:                   Gamma   Df Model:                            1
Link Function:          inverse_power   Scale:                         0.49076
Method:                          IRLS   Log-Likelihood:                -601.92
Date:                Sat, 20 Apr 2019   Deviance:                       56.351
Time:                        11:23:18   Pearson chi2:                     46.1
No. Iterations:                     7   Covariance Type:             nonrobust
===============================================================================
                  coef    std err          z      P>|z|      [0.025      0.975]
-------------------------------------------------------------------------------
Intercept       0.0037      0.000      8.701      0.000       0.003       0.005
Bicarbonate   2.78e-05   1.06e-05      2.634      0.008    7.11e-06    4.85e-05
===============================================================================
```

#### Omp-Pst2 porin – Bicarbonate – All statistic parameters:

```
                 Generalized Linear Model Regression Results                  
==============================================================================
Dep. Variable:                      Y   No. Observations:                   50
Model:                            GLM   Df Residuals:                       48
Model Family:                   Gamma   Df Model:                            1
Link Function:          inverse_power   Scale:                        0.016299
Method:                          IRLS   Log-Likelihood:                -246.36
Date:                Sat, 20 Apr 2019   Deviance:                      0.76802
Time:                        11:34:22   Pearson chi2:                    0.782
No. Iterations:                    18   Covariance Type:             nonrobust
=============================================================================================
                                coef    std err          z      P>|z|      [0.025      0.975]
---------------------------------------------------------------------------------------------
Intercept                     0.0013   3.89e-05     33.427      0.000       0.001       0.001
Bicarbonate                1.703e-05   1.17e-06     14.569      0.000    1.47e-05    1.93e-05
Concentration                 0.0013   3.89e-05     33.427      0.000       0.001       0.001
Bicarbonate:Concentration  1.703e-05   1.17e-06     14.569      0.000    1.47e-05    1.93e-05
=============================================================================================
```

#### Omp-Pst2 porin – Bicarbonate:

```
                 Generalized Linear Model Regression Results                  
==============================================================================
Dep. Variable:                      Y   No. Observations:                   50
Model:                            GLM   Df Residuals:                       48
Model Family:                   Gamma   Df Model:                            1
Link Function:          inverse_power   Scale:                        0.016299
Method:                          IRLS   Log-Likelihood:                -246.36
Date:                Sat, 20 Apr 2019   Deviance:                      0.76802
Time:                        11:35:15   Pearson chi2:                    0.782
No. Iterations:                     6   Covariance Type:             nonrobust
===============================================================================
                  coef    std err          z      P>|z|      [0.025      0.975]
-------------------------------------------------------------------------------
Intercept       0.0026   7.79e-05     33.427      0.000       0.002       0.003
Bicarbonate  3.405e-05   2.34e-06     14.569      0.000    2.95e-05    3.86e-05
===============================================================================
```

### 2-2 comparison (Tukey) – Ammonia condition

#### Omp-Pst1 porin

```
Multiple Comparison of Means - Tukey HSD,FWER=0.05
=================================================
group1 group2 meandiff   lower     upper   reject
-------------------------------------------------
  0     100   -226.57  -347.8389 -105.3011  True 
  0     500   -207.49  -328.7589  -86.2211  True 
  0     1000  -230.26  -351.5289 -108.9911  True 
 100    500    19.08   -120.9493  159.1093 False 
 100    1000   -3.69   -143.7193  136.3393 False 
```

```
 500    1000   -22.77  -162.7993  117.2593 False
```

#### Omp-Pst2 porin

```
Multiple Comparison of Means - Tukey HSD,FWER=0.05
===============================================
group1 group2 meandiff  lower    upper   reject
-----------------------------------------------
  0     100    98.33     2.99    193.67   True
  0     500    193.62   98.28    288.96   True
  0     1000   308.39  205.411  411.369   True
 100    500    95.29   -14.7991 205.3791 False
 100    1000   210.06  93.2928  326.8272  True
 500    1000   114.77  -1.9972  231.5372 False
```

#### 2-2 comparison (Tukey) – Bicarbonate condition

#### Omp-Pst1 porin

```
Multiple Comparison of Means - Tukey HSD,FWER=0.05
=================================================
group1 group2 meandiff   lower     upper   reject
-------------------------------------------------
  0      10    -57.92  -125.4145   9.5745  False
  0      25    -71.63   -126.739  -16.521   True
  0      50    -185.4   -240.509  -130.291  True
  0     100   -260.72   -315.829  -205.611  True
  10     25    -13.71   -81.2045  53.7845  False
  10     50   -127.48  -194.9745  -59.9855  True
  10    100    -202.8  -270.2945 -135.3055  True
  25     50   -113.77   -168.879  -58.661   True
  25    100   -189.09   -244.199  -133.981  True
  50    100    -75.32   -130.429  -20.211   True
```

#### Omp-Pst2 porin

```
Multiple Comparison of Means - Tukey HSD,FWER=0.05
=================================================
group1 group2 meandiff   lower     upper   reject
-------------------------------------------------
  0      10   -116.89  -151.1648  -82.6152  True
  0      25   -139.98  -174.2548 -105.7052  True
  0      50   -221.96  -256.2348 -187.6852  True
  0     100   -242.87  -277.1448 -208.5952  True
  10     25    -23.09   -57.3648  11.1848  False
  10     50   -105.07  -139.3448  -70.7952  True
  10    100   -125.98  -160.2548  -91.7052  True
  25     50    -81.98  -116.2548  -47.7052  True
  25    100   -102.89  -137.1648  -68.6152  True
  50    100    -20.91   -55.1848  13.3648   True
```

#### 2-2 comparison (Tukey) – pH condition

#### Omp-Pst1 porin

```
Multiple Comparison of Means - Tukey HSD,FWER=0.05
=================================================
group1 group2 meandiff   lower     upper   reject
-------------------------------------------------
  5      6     -110.6  -137.2876  -83.9124  True
  5      7     14.73    -11.9576  41.4176  False
  5      8    -192.42  -219.1076 -165.7324  True
  5      9    -165.36  -192.0476 -138.6724  True
  6      7     125.33   98.6424   152.0176  True
  6      8     -81.82  -108.5076  -55.1324  True
  6      9     -54.76   -81.4476  -28.0724  True
  7      8    -207.15  -233.8376 -180.4624  True
  7      9    -180.09  -206.7776 -153.4024  True
  8      9     27.06     0.3724   53.7476   True
```
